# Elevated plasma and urinary erythritol is a biomarker of excess simple carbohydrate intake in mice

**DOI:** 10.1101/2022.12.04.519026

**Authors:** Semira R. Ortiz, Martha S. Field

## Abstract

**Background:** Elevated serum erythritol is a predictive biomarker of diabetes and cardiovascular incidence and complications. Erythritol is synthesized endogenously from glucose, but little is known regarding the origin of elevated circulating erythritol *in vivo*.

**Objective:** *In vitro* evidence indicates that intracellular erythritol is elevated by high-glucose cell culture conditions and that final step of erythritol synthesis is catalyzed by the enzymes SORD and ADH1. The purpose of this study was to determine if dietary intake and/or diet-induced obesity (DIO) affect erythritol synthesis in mice, and if this relationship is modified by loss of the enzymes SORD or ADH1.

**Methods:** First, 8-week-old, male *Sord*^**+/+**^, *Sord*^-/-^, *Adh1*^+/+^, and *Adh1*^-/-^ mice were fed either low-fat diet (LFD) with 10% fat-derived calories or DIO high-fat diet (HFD) with 60% fat-derived calories for 8 weeks. Plasma and tissue erythritol were measured using GC-MS. Second, wild-type 8-week-old C57BL/6J mice were fed LFD or HFD with plain drinking water or 30% sucrose water for 8 weeks. Blood glucose and plasma and urinary erythritol were measured in non-fasted and fasted samples. Tissue erythritol was measured following sacrifice. Finally, *Sord*^**+/+**^ and *Sord*^-/-^ mice were fed LFD with 30% sucrose water for two weeks, then non-fasted plasma, urine, and tissue erythritol were quantified.

**Results:** Plasma and tissue erythritol were not impacted by loss of *Sord* or *Adh1* on LFD or HFD. In wild-type mice, consumption of 30% sucrose water significantly elevated plasma and urinary erythritol on both LFD and HFD compared to plain water. *Sord* genotype did not affect plasma or urinary erythritol in response to sucrose feeding, but *Sord*^-/-^ mice had reduced kidney erythritol content compared to wildtype littermates in response to sucrose.

**Conclusions:** Sucrose intake, not high-fat diet, elevates erythritol synthesis and excretion in mice. Loss of ADH1 or SORD does not significantly impact erythritol levels in mice.

## Introduction

Cardiometabolic diseases (such as diabetes, heart attack, and non-alcoholic fatty liver disease) begin to develop decades before clinical markers are apparent. The discovery and characterization of new biomarkers can facilitate early detection, and thus early intervention to prevent chronic disease progression. Serum erythritol is one biomarker with the potential to detect the metabolic dysregulation that precedes cardiometabolic diseases (1). Prospective cohort studies indicate that serum erythritol is elevated decades before the incidence of Type 2 Diabetes Mellitus and cardiovascular disease (2–8). Elevated erythritol has been consistently identified as a biomarker not only for future disease incidence, but also for worse outcomes in diagnosed patients (9–11).

Erythritol is a polyol traditionally thought of as a nonnutritive sweetener but was also recently found to be synthesized by humans through the pentose phosphate pathway (PPP) (4). Labelled erythritol appeared in plasma following the ingestion of universally labelled ^13^C-glucose, indicating that erythritol was synthesized from glucose (4). Ex vivo analysis of whole blood further indicated that erythritol is produced from erythrose-4-phosphate through the non-oxidative PPP (4). Little is known regarding the physiological role of erythritol in mammals.

Two mammalian enzymes have been identified that convert erythrose to erythritol: alcohol dehydrogenase 1 (ADH1) and sorbitol dehydrogenase (SORD) (12). ADH1 and SORD are homologous dehydrogenases (12). SORD is a strong candidate to catalyze erythritol synthesis in mammals. Knockdown of *SORD* in cell culture models reduces erythritol synthesis by 40% in high glucose conditions (12). In mice, tissues containing the most endogenous erythritol (the liver and kidney) are also the metabolic tissues in which SORD expression are highest (13). The impact of ADH1 expression on erythritol synthesis has not been explored. In human cells, erythritol synthesis also exhibits a dose-response to the amount of glucose provided in culture media, suggesting that erythritol may respond to nutrient excess (14).

There have been no studies on the factors that contribute to endogenous erythritol synthesis *in vivo* in mammals. The purpose of this work was to determine the role of the enzymes SORD and ADH1 in erythritol synthesis and how erythritol levels are impacted by diet *in vivo*. This is the first study to report that erythritol synthesis and excretion is elevated in response to a high-sucrose diet.

## Methods

### Generation of the *Sord*^-/-^ mice

We used crispr.mit.edu to select the guide RNA sequence with minimal off-target effects, targeting exon 4 of *Sord* (guide sequence: 5’-AGAAGAAGATAGTCGCGCTC-3’). Template DNA was generated by PCR using the forward primer: 5’-GAAATTAATACGACTCACTATAGGAGAAGAAGATAGTCGGCGTCGTTTTAGAGCTA

GAAATAGC-3’ and reverse primer: 5’-GCACCGACTCGGTGCCACTTTTTCAAGTTGATAACGGACTAGCCTTATTTTAACTTGC TATTTCTAGCTCTAAAAC-3’. Two identical 50 μL reactions were prepared consisting of 25 μL GoTaq DNA Polymerase (Promega), 0.5 μL forward primer, 0.5 μL reverse primer, and 24 μL nuclease-free water. DNA was denatured at 94°C for 5 minutes, followed by 40 cycles of 30 second denaturation at 94°C, 30 second annealing at 58°C, 20 second extension at 72°C. Final elongation was performed at 72°C for 5 minutes. PCR product was pooled, and the DNA amplicon was purified using the MinElute PCR Purification Kit (Qiagen) per manufacturer’s instructions. DNA quality was checked by NanoDrop and agarose gel.

*In vitro* transcription was completed using the MegaShortscript T7 Transcription kit (ThermoFisher) according to the manufacturer’s protocol. Transcription incubation was performed overnight at 37°C in a dry incubator. RNA was then purified using the MEGAClear Transcription Clean-Up Kit (ThermoFisher) per manufacturer protocol. RNA quantity was determined by Qubit. Quality was verified by denaturing 1 μL RNA, 1 μL formaldehyde loading dye (Ambion), and 8 μL nuclease free water in a thermocycler at 65°C for 10 minutes. The denatured RNA was run on an agarose gel to check for the presence of a single sgRNA species. RNA was stored at -80°C until microinjection.

Embryos were isolated from 15 C57BL/6J donor females. Pure sgRNA and Cas9 mRNA were microinjected into the pronucleus and cytoplasm of 203 1 cell embryos. Of the 1 cell embryos, 174 advanced to the 2-cell stage and were transferred equally to 6 pseudo-pregnant recipient female mice. 50 founder (F0) pups were born.

DNA was isolated from tail snips with the High Pure PCR Template Preparation Kit (Roche) per the manufacturer’s instructions. The 400 base pair region surrounding the sgRNA target was then amplified by PCR (forward primer: 5’-CCCAGAGAGGAGGCTGTAGA-3’; reverse primer: 5’-AAAGGCCTCCCAGGGGTTAT-3’) with GoTaq DNA Polymerase (Promega). The resulting PCR product was cloned into the pCR 4-TOPO vector using the TOPO TA Cloning Kit for Sequencing (Invitrogen). Briefly, 4 μL PCR product, 1 μL Salt Solution, and 1 μL pCR 4-TOPO Vector were combined and incubated for 5 minutes at room temperature. The TOPO cloning reaction was then transformed into One Shot TOP10 Chemically Competent *E. coli* and plated on 50 μg/mL kanamycin LB plates. After incubation overnight at 37°C, 4 colonies per mouse were picked and cultured overnight in 4 mL LB medium with 50 μg/mL kanamycin. Vector DNA was purified using the Zyppy Plasmid Miniprep Kit (Zymo Research) according to the manufacturer’s protocol. Plasmids were analyzed by Sanger sequencing to detect mutations in the *Sord* gene (M13 Forward (-20) primer: 5’-GTAAAACGACGGCCAG - 3’).

F0 males with a confirmed mutation in the *Sord* gene were mated with C57BL/6J females to obtain heterozygous F1 pups. *Sord* deletion was confirmed by a 50% reduction in liver SORD protein in F1 *Sord*^**+/-**^ mice, measured by western blot analysis. *Sord*^**+/-**^ male and female mice were mated to produce *Sord*^**+/+**^ (wildtype, WT) and *Sord*^-/-^ (knockout, KO) mice.

### Animal dietary treatments and tissue collection

All mice were maintained under specific-pathogen-free conditions in accordance with standard of use protocols and animal welfare regulations. All study protocols were approved by the Institutional Animal Care and Use Committee of Cornell University. All mice were housed individually in environmentally controlled conditions (12 hour light/12 hour dark cycle). *Adh1* mice have been previously described and were backcrossed 10 generations to a C57Bl/6J background (15). At 8 weeks of age, male *Sord*^**+/+**^, *Sord*^-/-^, *Adh1*^+/+^, and *Adh1*^-/-^ mice were randomly assigned to one of two diets: low-fat diet (LFD) with 10% fat-derived calories or diet-induced obese (DIO) high-fat diet (HFD) with 60% fat-derived calories. Diets were based on the AIN-93G Purified Rodent Diet (Dyets Inc., Bethlehem PA, DYET#’s 104783 and 103651). Diet compositions are provided in **Supplementary Table 1 (LFD) and 2 (HFD)**. Food and water were provided *ad libitum* for 8 weeks. Food intake and body weight were measured twice weekly. Food intake was determined by subtracting the weight of food remaining in the hopper from the weight of food that was supplied. Body composition was measured by NMR after 2, 5, and 8 weeks of treatment using a Bruker Minispec LF65 according to the manufacturer’s protocols. Body composition measurements included free fluid, lean, and fat mass.

To measure the impact of sugar intake on erythritol synthesis, additional 8-week-old male C57BL/6J mice were randomly assigned to one of four diets for eight weeks. Diets included low-fat diet with plain drinking water (LFD), high-fat diet with plain drinking water (HFD), LFD with 30% sucrose in drinking water (LFD+30% Sucrose) and HFD with 30% sucrose in drinking water (HFD+30% Sucrose). Body weight, food intake, and body composition were assessed as described above. Caloric intake from water was calculated from milliliters of 30% sucrose consumed.

Following dietary treatment, all mice were killed by carbon dioxide asphyxiation and cervical dislocation. Plasma and tissues (adipose, liver, kidney, quadriceps) were harvested and snap-frozen in liquid nitrogen, followed by storage at -80°C for use in later applications.

### Intraperitoneal glucose tolerance testing

Mice were fasted for 5 hours prior to intraperitoneal glucose tolerance testing (IPGTT). Mice were injected intraperitoneally with 1.5 mg glucose/g body mass. Blood glucose was measured at 15, 30, 60, 90, and 120 minutes following glucose injection. All blood samples were collected from a single nick in the tail vein. Blood glucose was measured via a drop of blood applied to a hand-held glucometer (OneTouch). The area under the curve was calculated using Prism software.

### Collection of plasma and urine

Blood was collected from a nick in the tail vein into an EDTA-coated microvette tube (Sarstedt). Whole blood was centrifuged for 10 minutes at 2,000 x g and 4C, then plasma was transferred to an Eppendorf tube and stored at -80C for later analysis of plasma metabolites. Urine samples were collected as previously described (16). Fasted samples were collected following a 5-hour daytime fast.

### Isolation and measurement of polar metabolites by GC-MS

Polar metabolites were isolated from mouse plasma, tissues, and urine as described previously (13). Urine was diluted 1 part to 2 parts Milli-Q water prior to isolation with extraction fluid. To account for urine concentration, creatinine was measured using the Creatinine (urinary) Colorimetric Assay Kit per the manufacturer’s protocol (Cayman Chemical).

Dried metabolites were then derivatized and analyzed by GC-MS as previously described (13). In SIM mode, mass spectra of m/z 217, m/z 307, and m/z 320 were acquired from 8-9 min, m/z 218, m/z 320, and m/z 423 were acquired from 10-11 min, and m/z 319, m/z 331, and m/z 421 were acquired from 12-13 minutes. Erythritol, sorbitol, and ^13^C_1_-ribitol peaks were selected from GC-MS chromatograms based on the retention time of their respective standards. Absolute intensities of erythritol (m/z 217), sorbitol (m/z 319), and ^13^C_1_-ribitol (m/z 218) were recorded. The ratio of the absolute intensity of erythritol or sorbitol to that of ribitol (relative intensity) was used to determine plasma erythritol concentration. Tissue samples were normalized by dividing the relative metabolite intensity by tissue mass in grams. Urine samples were normalized by dividing the relative metabolite intensity by the sample creatinine concentration.

### Western blot analysis

Frozen tissue samples were homogenized in lysis buffer containing 15% NaCl, 5 mM EDTA, pH 8, 1% Triton X100, 10 mM Tris-Cl, 5 mM DTT, and 10 µl/mL protease inhibitor cocktail (Sigma Aldrich). Protein concentration was determined by Lowry assay (17). Equal amounts of protein (25-50ug) were denatured by heating with 6X Laemelli buffer for 5-10 min at 95 °C. Samples were then loaded onto a 10% SDS-PAGE gel and electrophoresed. Protein was transferred by electrophoresis to an Immobilon-P PVDF membrane (Millipore Corp.).

The membrane was blocked in 5% non-fat milk overnight at 4°C, then incubated with primary antibodies for 1 hour at room temperature. Primary antibodies included sorbitol dehydrogenase (1:2,000, Proteintech), alcohol dehydrogenase 1, lamin B1, transketolase, and alpha-tubulin (1:1,000, Cell Signaling Technology). Secondary anti-rabbit antibody (1:100,000, ThermoFisher) was applied to the membrane and incubated for 1 hour at room temperature. Protein was detected using a Protein Simple FluorChem E with Clarity Western ECL Substrate (Bio-Rad). Band intensity was measured using ImageJ (NIH).

### Statistical analysis

All statistical analyses were conducted using GraphPad Prism 9 (Graphpad Software Inc). No blinding to treatment group was performed. Differences between two groups (sucrose pilot and sucrose exposure in SORD animals) were analyzed using two-sided unpaired t-tests. Two-way ANOVAs were used for analysis of interactions and main effects (genotype and diet or dietary fat and dietary sucrose). Sidak’s multiple comparisons test was used as post hoc analysis for ANOVA tests to determine differences between groups. The difference in macronutrient intake from carbohydrates was analyzed using one-way Welch’s ANOVAs to correct for unequal standard deviations. All tests were performed at the 95% confidence level (α = 0.05) and groups were considered significantly different when *p ≤0*.*05*. No criteria were set for animal exclusion *a priori*, and no experimental data points or animals were excluded from analysis.

## Results

### *Sord* knockout does not impact erythritol synthesis in mice

Loss of SORD protein was confirmed in liver and kidney by western blot **(Fig. 1A and 1B)**. We found no difference in body weight or caloric intake between SORD WT and KO animals on either LFD or HFD **(Fig. S1)**. After 8 weeks of dietary treatment, there was a main effect of *Sord* genotype on body fat percentage, however, no significant differences were detected in post-hoc analysis **(Fig. S2A, genotype effect p<0.05)**. There was also no *Sord* genotype-driven difference in adipose depot weight, regardless of diet (Fig. S2B and S2C). There was no significant effect of *Sord* genotype on glucose tolerance area under the curve **(Fig. S3)**.

**Figure 1.**
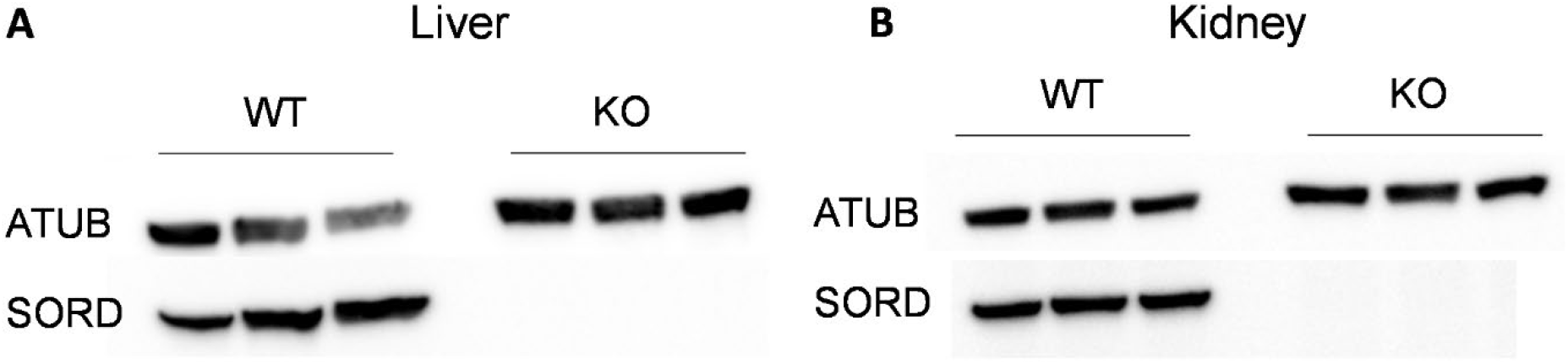
Single base deletion in exon 4 eliminates liver and kidney SORD protein. SORD protein is absent in A) liver and B) kidney of 8-week-old KO mice. Data points represent tissues harvested from individual mice (n=3). ATUB: alpha tubulin; KO: knockout; SORD: sorbitol dehydrogenase; WT: wildtype.

Fasted plasma erythritol was not modified by loss of SORD after 2 or 8 weeks of dietary treatment **(Fig. 2A and 2C)**. At 5 weeks, there was a significant main effect of genotype on fasting plasma erythritol, but no differences were detected in specific pairwise comparisons (Fig. 2B, ANOVA main effect of genotype p=0.01). We found no effect of diet on plasma erythritol at any time point (Fig. 2A-2C). There was also no effect of SORD loss on liver or kidney erythritol content **(Fig 3A, 3B)**. Unexpectedly, wild-type animals fed HFD had significantly less erythritol in the liver and kidneys (Fig. 3A, p<0.05 and 3B, p<0.01). This effect was not observed in KO mice.

**Figure 2.**
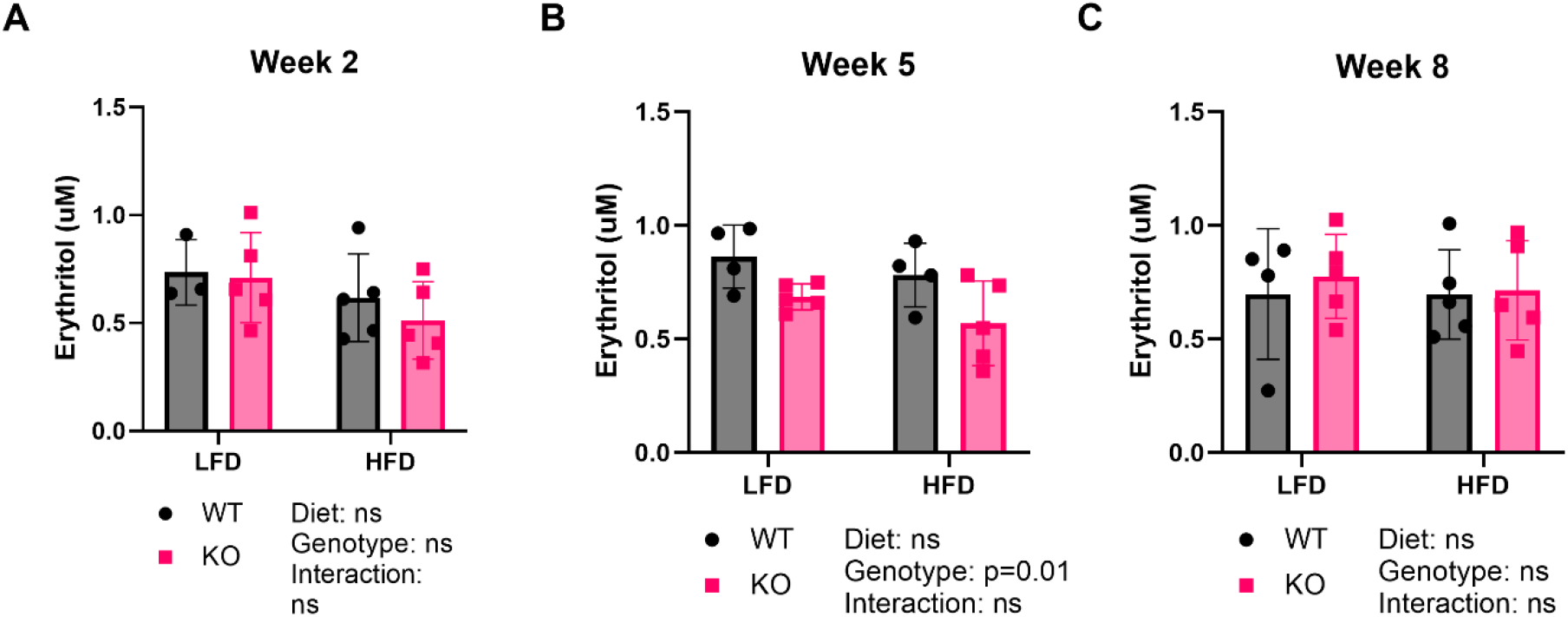
Loss of SORD does not affect plasma erythritol in mice fed LFD or HFD. Plasma erythritol in SORD WT and KO mice following A) 2 weeks, B) 5 weeks, and C) 8 weeks of treatment with LFD or HFD. Data presented as mean ± SD. HFD: high-fat diet; KO: knockout; LFD: low-fat diet; WT: wildtype.

**Figure 3.**
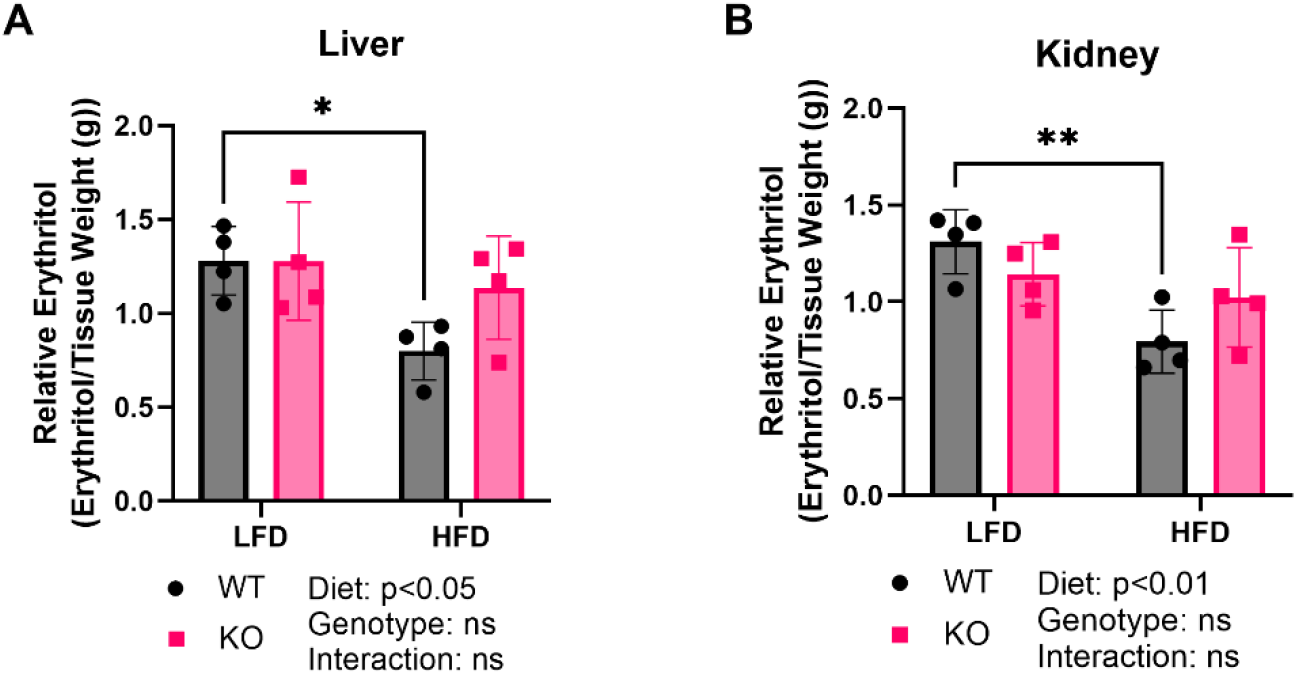
Loss of SORD does not affect tissue erythritol in mice fed LFD or HFD. Relative erythritol content of A) liver and B) kidney of SORD WT and KO mice after 8 weeks on experimental diets. Data presented as mean ± SD. *p<0.05, **p<0.01. HFD: high-fat diet; KO: knockout; LFD: low-fat diet; WT: wildtype.

To determine if SORD deletion results in sorbitol accumulation, we assessed plasma and tissue sorbitol levels. Indeed, we found significantly elevated plasma sorbitol in KO compared to WT littermates **(Fig. S4A, p<0.0001)**. Sorbitol accumulation was further exacerbated by HFD in SORD KO mice compared to LFD (Fig. S4A, p<0.05). Liver sorbitol was also significantly elevated in SORD KO mice on LFD and HFD (Fig. S4B, p<0.05 and p<0.001 respectively). There was no difference in kidney sorbitol (Fig. S4C).

To assess if the effect of SORD deletion on erythritol synthesis is blunted by a compensatory increase in ADH1 protein levels, we quantified liver and kidney ADH1 protein. We found no increase in ADH1 levels in KO compared to WT animals **(Fig. 4A and 4B)**. We also assessed expression of TKT, an enzyme in the non-oxidative pentose phosphate pathway that has been shown to regulate erythritol synthesis in cultured cells (14). TKT expression was reduced in the kidney of mice fed HFD compared to LFD (Fig. 4B, p<0.0001 and p<0.001 in WT and KO mice, respectively), which may explain the observed reduction in kidney erythritol in WT animals fed HFD (Fig 3B).

**Figure 4.**
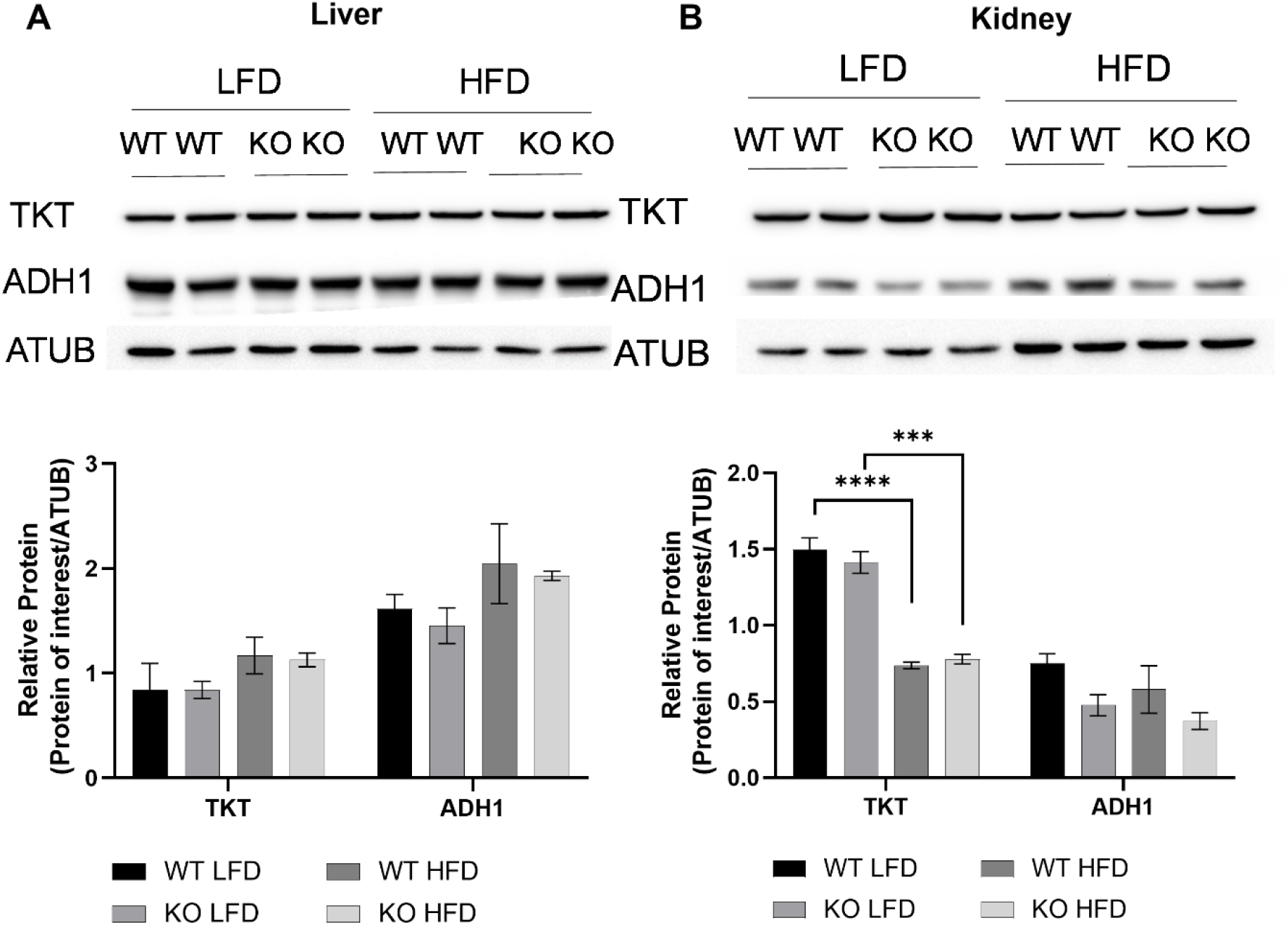
*Sord* genotype does not impact TKT or ADH1 levels in liver and kidney. Representative western blot and densitometry quantification of enzymes TKT and ADH1 in A) liver and B) kidney of SORD WT and KO mice after 8 weeks of dietary treatment. Data points represent tissue harvested from individual mice (n=2) and quantification is presented as mean ± SD. Differences between groups are analyzed by two-way ANOVA. *p<0.05, **p<0.01, ***p<0.001, ****p<0.0001. ADH1: alcohol dehydrogenase 1; ATUB: alpha tubulin; HFD: high-fat diet; KO: knockout; LFD: low-fat diet; SORD: sorbitol dehydrogenase; TKT: transketolase; WT: wildtype.

### *Adh1* knockout has no effect on plasma or tissue erythritol

We next assessed the impact of loss of ADH1 on erythritol synthesis. ADH1 KO animals displayed normal body weight, caloric intake, body composition, and glucose tolerance compared to WT littermates **(Fig. S5-S7)**. Loss of ADH1 did not impact fasting plasma erythritol levels at any time point **(Fig. 5)**. Consistent with results in SORD animals, we also found no differences in plasma erythritol between diets (Fig. 5). Similarly, there was no effect of genotype or diet on tissue erythritol **(Fig. 6A and 6B)**. We did not observe any differences in SORD or TKT expression in the liver or kidney between ADH1 WT and KO groups, regardless of diet **(Fig. 7A and 7B)**.

**Figure 5.**
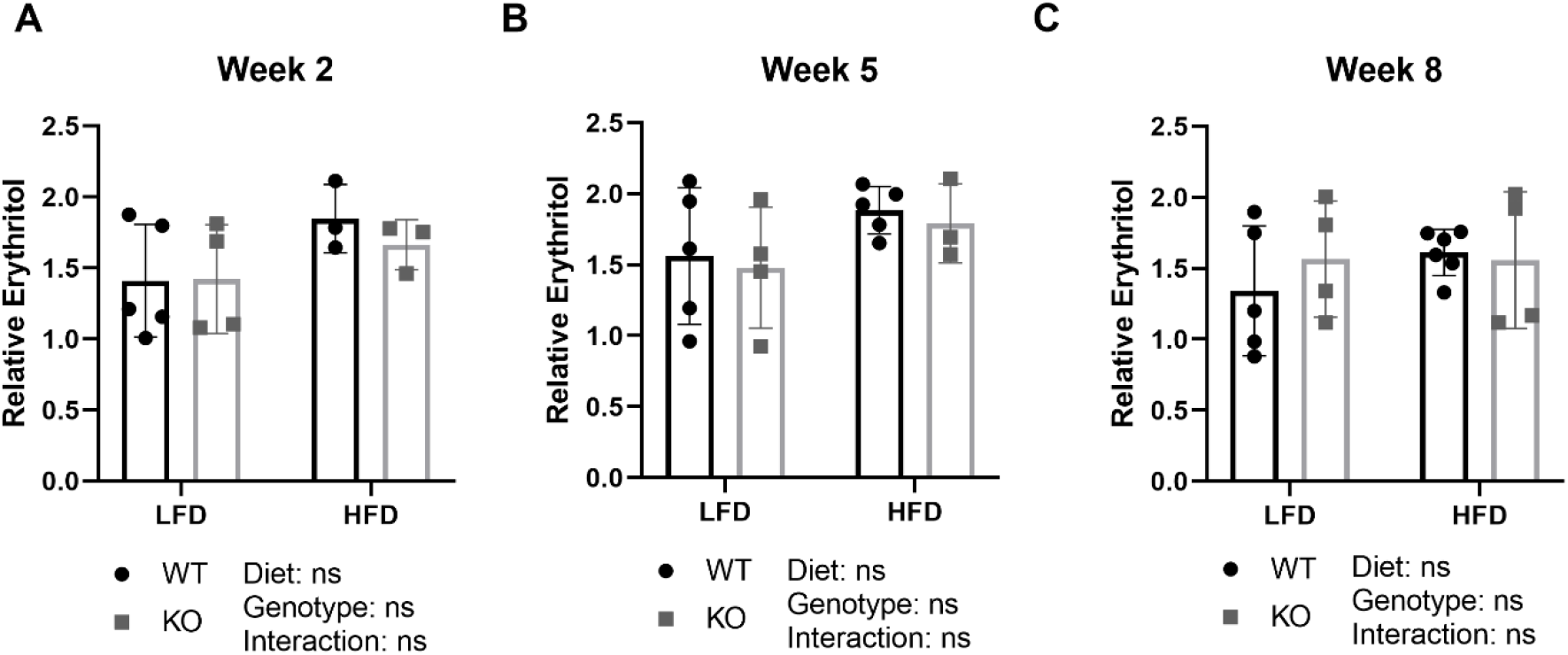
Plasma erythritol is not impacted by *Adh1* knockout. Plasma erythritol in ADH1 WT and KO mice following A) 2 weeks, B) 5 weeks, and C) 8 weeks of treatment with LFD or HFD. Data presented as mean ± SD. HFD: high-fat diet; KO: knockout; LFD: low-fat diet; WT: wildtype.

**Figure 6.**
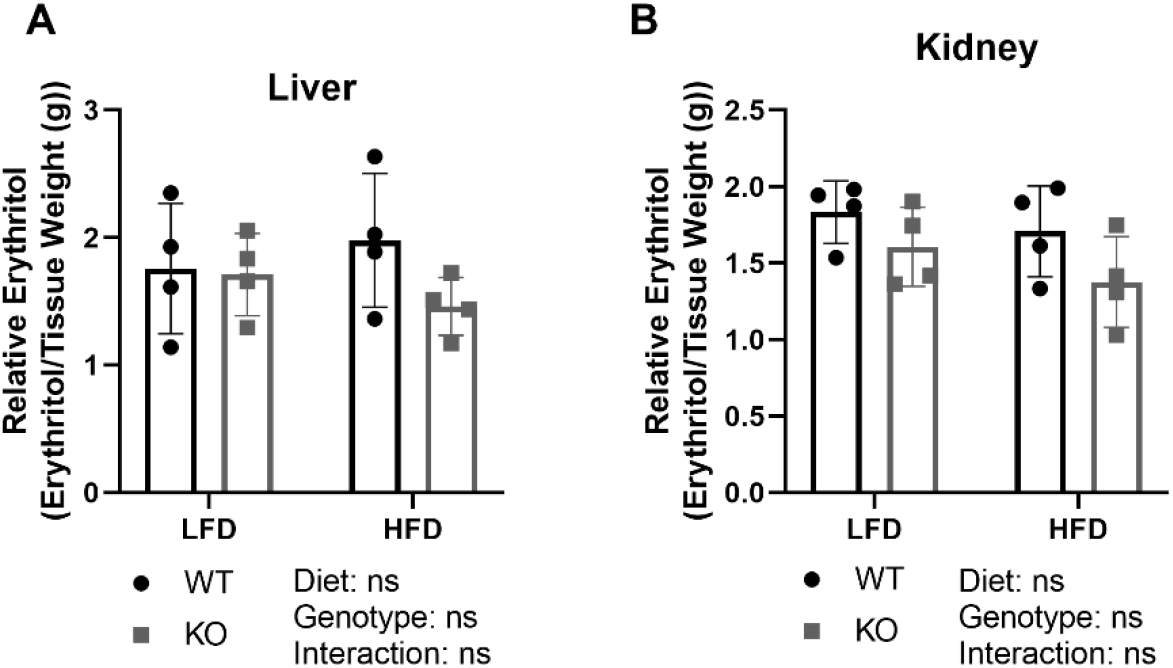
*Adh1* deletion does not significantly decrease liver or kidney erythritol. Relative erythritol content of A) liver and B) kidney of ADH1 WT and KO mice after 8 weeks on experimental diets. Data presented as mean ± SD. HFD: high-fat diet; KO: knockout; LFD: low-fat diet; WT: wildtype.

**Figure 7.**
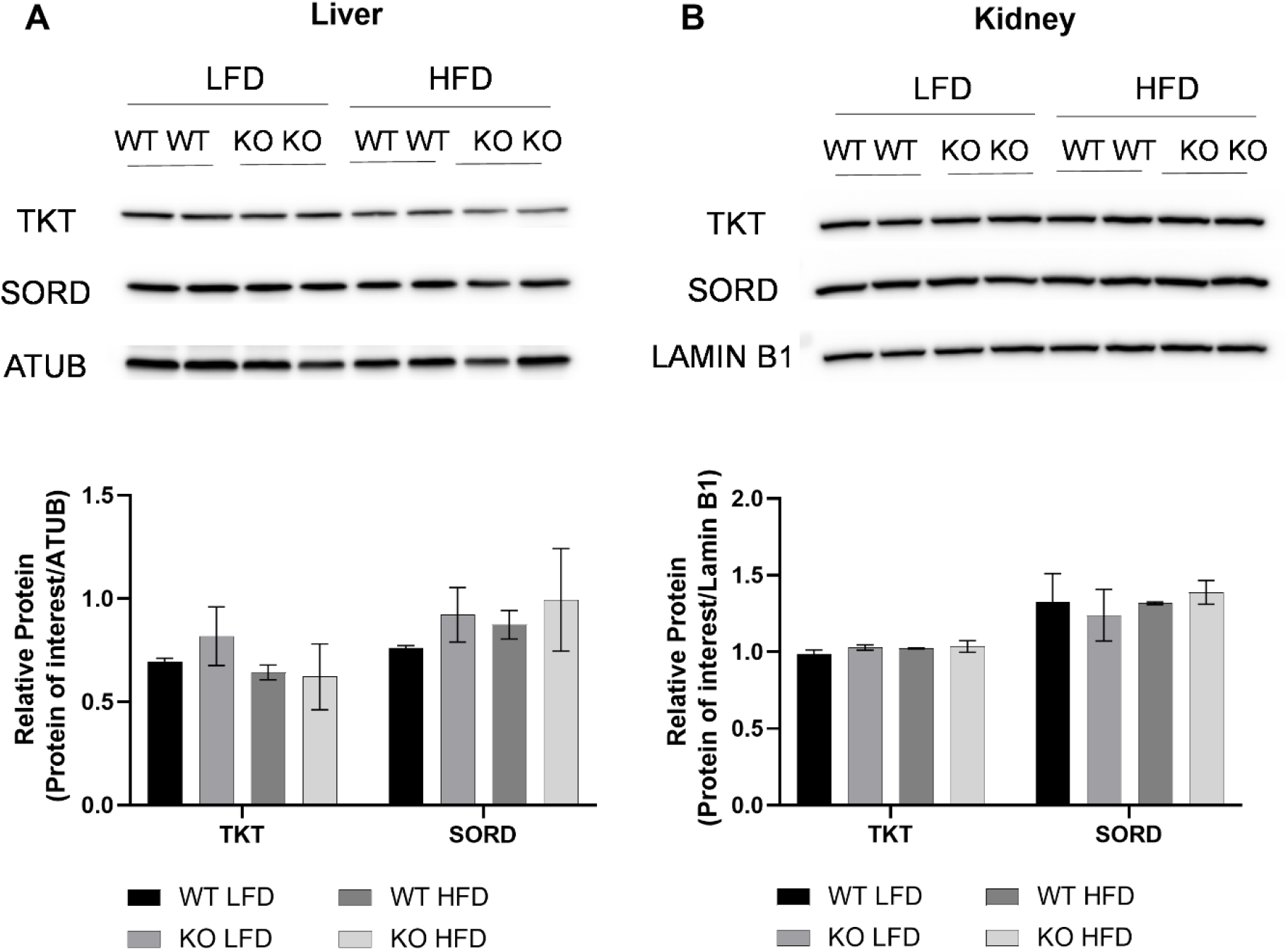
*Adh1* genotype does not impact TKT or SORD expression in liver and kidney. Representative western blot and densitometry quantification of enzymes TKT and SORD in A) liver and B) kidney of ADH1 WT and KO mice after 8 weeks of dietary treatment. Data points represent tissue harvested from individual mice (n=2) and quantification is presented as mean ± SD. Differences between groups are analyzed by two-way ANOVA. ADH1: alcohol dehydrogenase 1; ATUB: alpha tubulin; HFD: high-fat diet; KO: knockout; LFD: low-fat diet; SORD: sorbitol dehydrogenase; TKT: transketolase; WT: wildtype.

### Erythritol synthesis is responsive to sucrose consumption even in the absence of changes of body weight or fasting blood glucose

In SORD and ADH1 animal models, we observed no effect of HFD on circulating erythritol, despite significant body weight gain and impaired glucose tolerance in HFD fed mice. This suggests that erythritol synthesis in mice is not sensitive to hyperglycemia. Erythritol synthesis, then, may respond to diet-induced increases in glucose availability. To test this hypothesis, we provided plain drinking water or 30% sucrose solution to C57BL/6J mice fed HFD for two weeks.

30% sucrose for two weeks did not affect body weight, although total caloric intake was significantly elevated **(Fig. S8A and S8B, p<0.01)**. Fasting blood glucose was not affected by 30% sucrose (S8C). There was no difference in fasted plasma erythritol between groups **(Fig. S9A)**. Surprisingly, we found a 50% increase in non-fasted urine erythritol in mice fed 30% sucrose in drinking water (Fig. S9B p<0.05).

To evaluate the response of erythritol synthesis to diet more comprehensively, we utilized the addition of 30% sucrose in drinking water to LFD or HFD for eight weeks. As expected, mice fed HFD gained significantly more body weight over the course of 8 weeks compared to LFD **(Fig. 8A)**. This was consistent in both HFD with water and HFD with 30% sucrose (Fig. 8A, p<0.01 and p<0.001 respectively for effect of dietary fat). HFD also significantly increased body fat percentage compared to respective LFD controls (Fig. 8B, p<0.0001). There was a main effect of sucrose on body fat percentage, however, no significant differences were detected in pairwise comparisons (Fig. 8B, ANOVA main effect of sucrose p<0.05). Sucrose water significantly increased total caloric intake in mice fed LFD and HFD (Fig. 8C, p<0.01). This increase in total calories in mice fed sucrose water resulted from a 2-fold (LFD) and 4-fold (HFD) increase in carbohydrate intake compared to plain water controls (Fig. 8D, one-way ANOVA, p<0.0001 and p<0.001 respectively).

**Figure 8.**
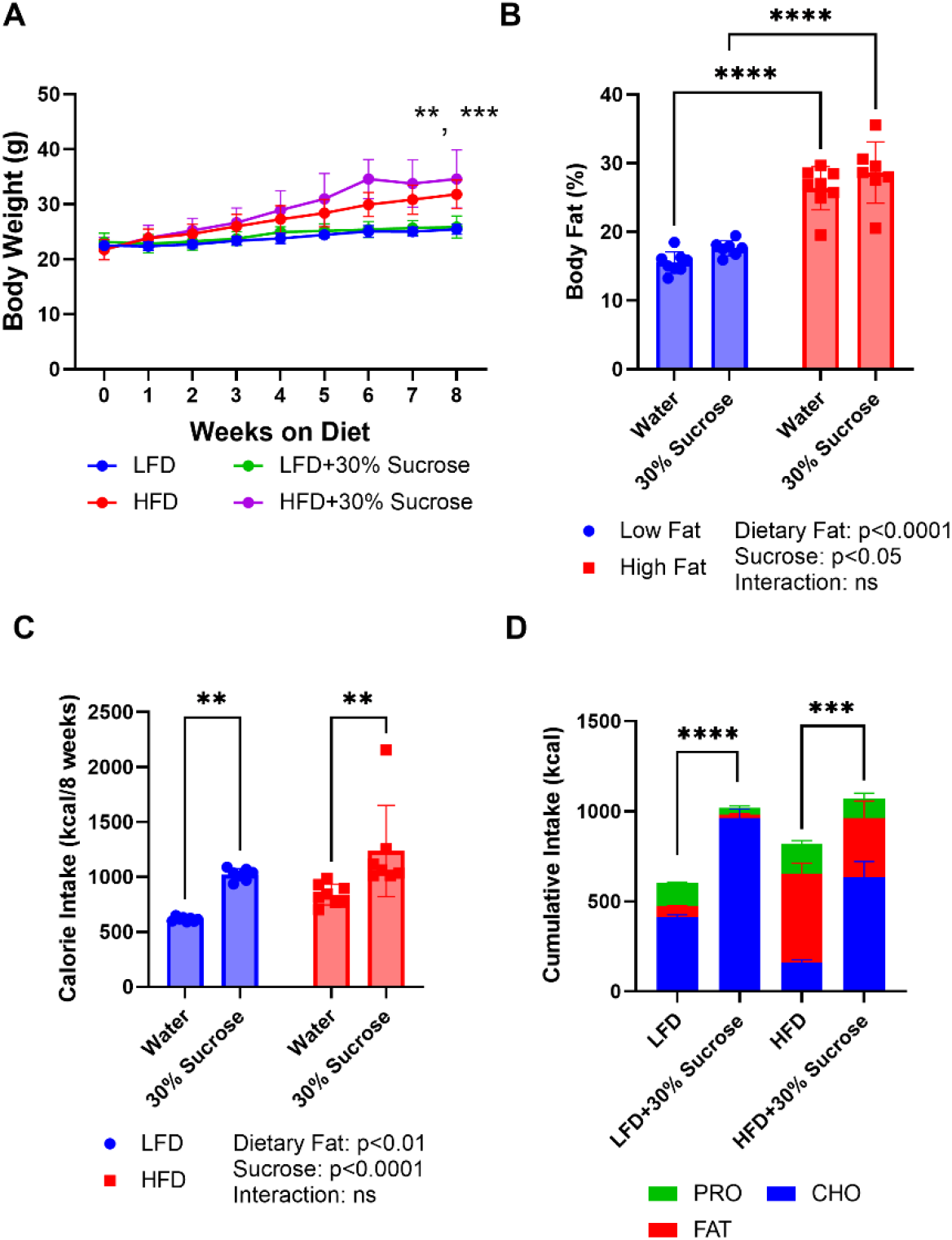
Sucrose in drinking water significantly increased caloric intake, body weight and fat percentage. A) Weekly body weight in grams, asterisks indicate p-value comparing LFD to HFD and LFD + 30% sucrose to HFD + 30% sucrose, respectively, at the 8-week timepoint. B) Body fat percentage following 8 weeks of dietary treatment measured by NMR. C) Total cumulative caloric intake in kilocalories (kcal). D) Proportion of total caloric intake from protein, carbohydrates, and fat in each experimental diet. Asterisks represent results of a one-way Welch’s ANOVA comparing cumulative intake from carbohydrates. Data are expressed as mean ± SD. **p<0.01, ***p<0.001, ****p<0.0001. CHO: carbohydrate; HFD: high-fat diet; LFD: low-fat diet; PRO: protein.

Dietary fat, but not sucrose water, contributed to changes in random and fasting blood glucose **(Fig. 9A-9D)**. At 5 weeks of dietary treatment mice fed HFD with 30% sucrose had higher random blood glucose levels than LFD with 30% sucrose (Fig. 9B, p<0.05). Fasting blood glucose was significantly higher in mice fed HFD and HFD with sucrose water compared to respective LFD controls (Fig. 9D, p<0.05). There was no effect of sucrose water on fasting glucose or random glucose at any timepoint (Fig. 9A-9D).

**Figure 9.**
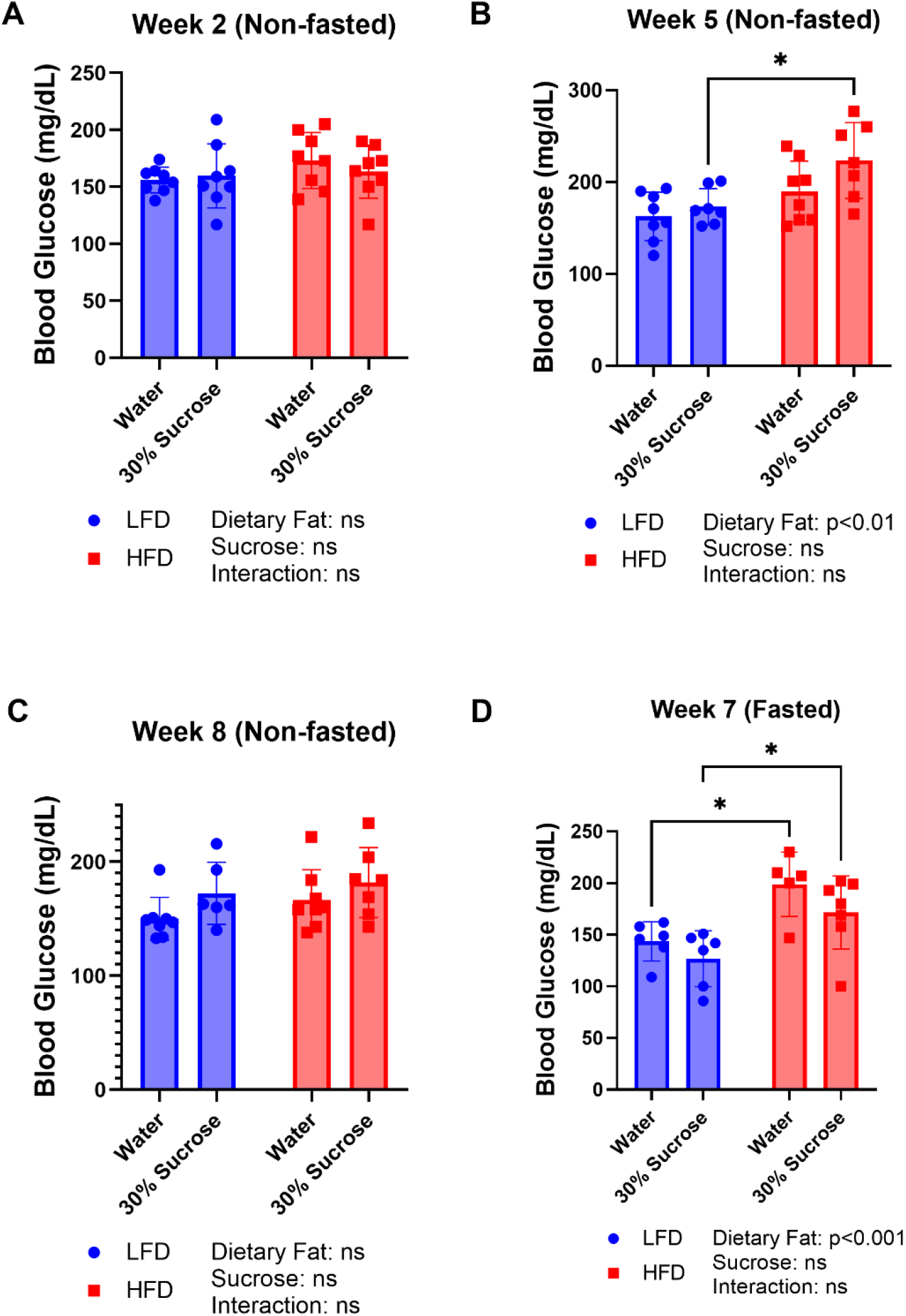
Sucrose in drinking water did not significantly impact fed or fasted blood glucose. Blood glucose in non-fasted mice at A) 2 weeks, B) 5 weeks, and C) 8 weeks of diet treatment. D) Blood glucose in fasted mice after 7 weeks of dietary treatment. Data are presented as mean ± SD. *p<0.05. HFD: high-fat diet; LFD: low-fat diet.

### Plasma and urine erythritol are elevated in response to sucrose water

After two weeks on experimental diets, sucrose in drinking water significantly increased non-fasted plasma erythritol on LFD (4.5-fold) and HFD (2.6-fold) compared to water controls **(Fig. 10A, p<0.0001 and p<0.05 respectively)**. Additionally, there was a significant interaction between dietary fat and sucrose (Fig. 10A, ANOVA interaction p<0.05). Mice consuming LFD with 30% sucrose had over 60% higher plasma erythritol compared to HFD with 30% sucrose (Fig. 10A, p<0.01). The sucrose-induced increase in plasma erythritol was consistent at 2, 5 and 8 weeks of dietary treatment in non-fasted samples (Fig. 10A, 10B, and 10C). Following a 5-hour fast, there were no differences in plasma erythritol between any of the 4 experimental diets at the 7 week timepoint (Fig. 10D).

**Figure 10.**
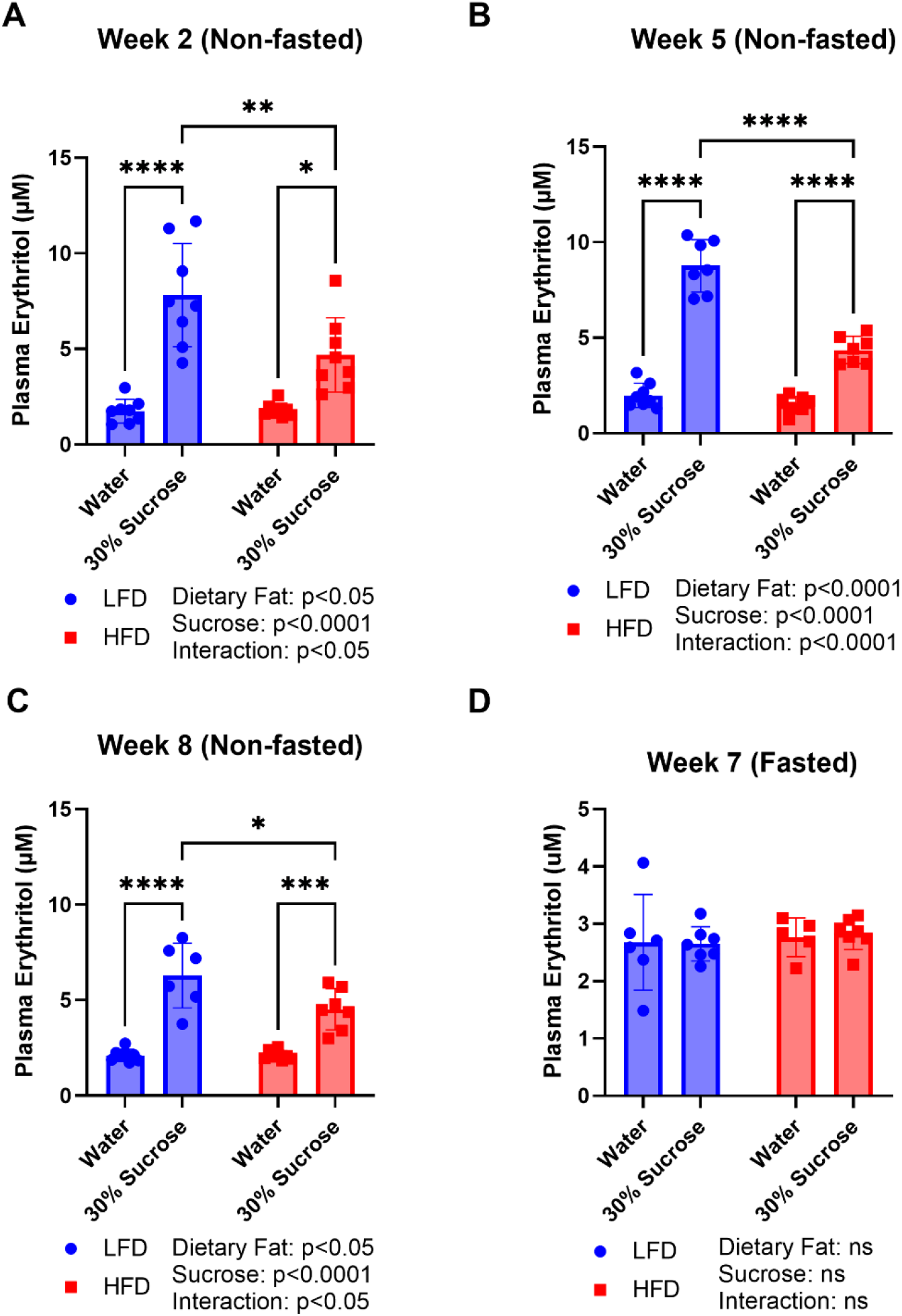
Plasma erythritol is significantly elevated by sucrose in fed, but not fasted mice. Plasma erythritol in non-fasted mice at A) 2 weeks, B) 5 weeks, and C) 8 weeks of diet treatment. D) Plasma erythritol in fasted mice after 7 weeks of dietary treatment. Data are presented as mean ± SD. *p<0.05, **p<0.01, ***p<0.001, ****p<0.0001. HFD: high-fat diet; LFD: low-fat diet.

Urine erythritol levels paralleled plasma erythritol levels. After two weeks exposure to experimental diets, there was a significant interaction between the effect of dietary fat and sucrose on non-fasted urine erythritol **(Fig. 11A, ANOVA interaction p<0.05)**. The LFD with 30% sucrose group was significantly higher than both LFD controls and HFD with 30% sucrose (Fig. 11A, p<0.0001 and p<0.01). After 5 weeks and 8 weeks, both LFD with 30% sucrose and HFD with 30% sucrose excreted significantly more erythritol than their respective controls (Fig. 11B and 11C). Consistent with plasma erythritol, there were no differences in fasted urine erythritol content (Fig. 11D).

**Figure 11.**
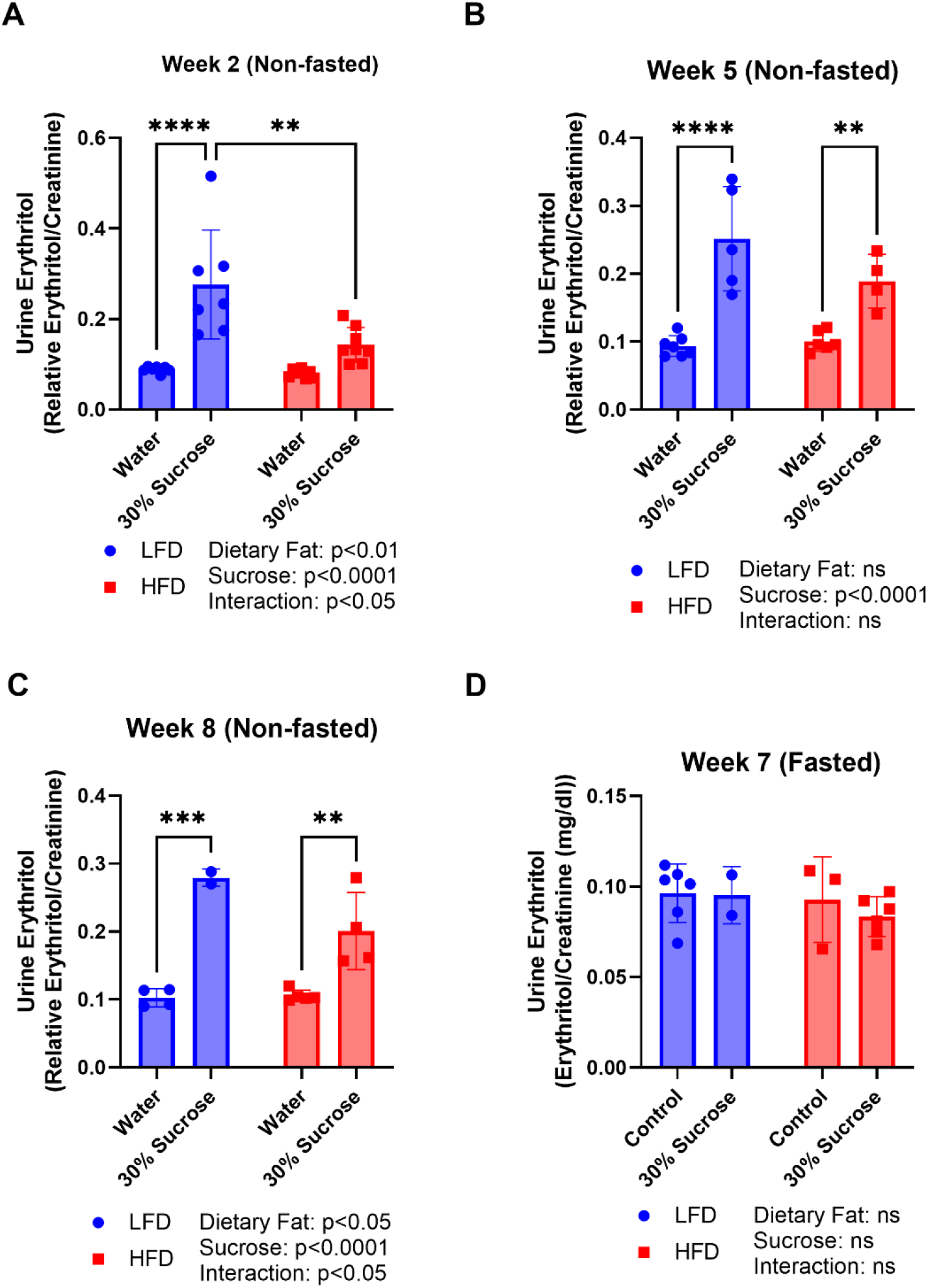
Urinary erythritol is elevated by sucrose in drinking water in fed mice. Relative urinary erythritol in non-fasted mice at A) 2 weeks, B) 5 weeks, and C) 8 weeks of diet treatment. D) Relative urinary erythritol in fasted mice after 7 weeks of dietary treatment. Erythritol was normalized to urinary creatinine content. Data are presented as mean ± SD. **p<0.01, ***p<0.001, ****p<0.0001. HFD: high-fat diet; LFD: low-fat diet.

### Plasma and urine sorbitol are increased by sucrose water

We also assessed plasma and urine levels of sorbitol to determine if additional endogenous polyols exhibit the same response to sucrose consumption. The effect of diet on plasma sorbitol levels varied across the 4 measured timepoints **(Fig S10A-S10D)**. At 2 weeks, there were significant main effects of fat and sucrose on plasma sorbitol, but no differences were detected in pairwise comparisons (Fig. S10A, ANOVA main effect of dietary fat p<0.05, main effect of sucrose p<0.05). At 5 weeks, sucrose water significantly elevated plasma sorbitol compared to water controls on LFD and HFD (Fig. S10B, p<0.05). After 8 weeks, only mice fed LFD with 30% sucrose had elevated plasma sorbitol compared to LFD mice (Fig. S10C, p<0.001). In fasted mice, plasma sorbitol was higher in HFD mice compared to LFD mice with drinking water (Fig. S10D, p<0.05).

Urine sorbitol was elevated at 2 weeks in the LFD with 30% sucrose group compared to both LFD and HFD with 30% sucrose **(Fig. S11A, p<0.01 and p<0.05)**. After 5 weeks, both sucrose-fed groups had significantly higher urine sorbitol compared to water controls (Fig. S11B, p<0.05). There was a main effect of sucrose on urine sorbitol, but no significant pairwise comparisons of urine sorbitol after 8 weeks (Fig. S11C, ANOVA main effect of sucrose p<0.05). There were no differences in fasting urine sorbitol (Fig. S11D).

### The effect of sucrose water on tissue erythritol is tissue-dependent

The kidneys, liver, and quadriceps have previously been shown to synthesize erythritol (13). Under low- and high-fat dietary conditions, the kidneys contain the highest levels of erythritol per gram tissue, followed closely by the liver (13). We found no significant difference in liver erythritol in response to sucrose water exposure **(Fig. 12A)**. In the kidney, sucrose water on both LFD and HFD caused a 3-fold increase in erythritol compared to respective controls (Fig. 12B, p<0.01). Unexpectedly, sucrose water also elevated quadriceps erythritol by more than 3-fold on both LFD and 2-fold on HFD (Fig. 12C, p<0.001).

**Figure 12.**
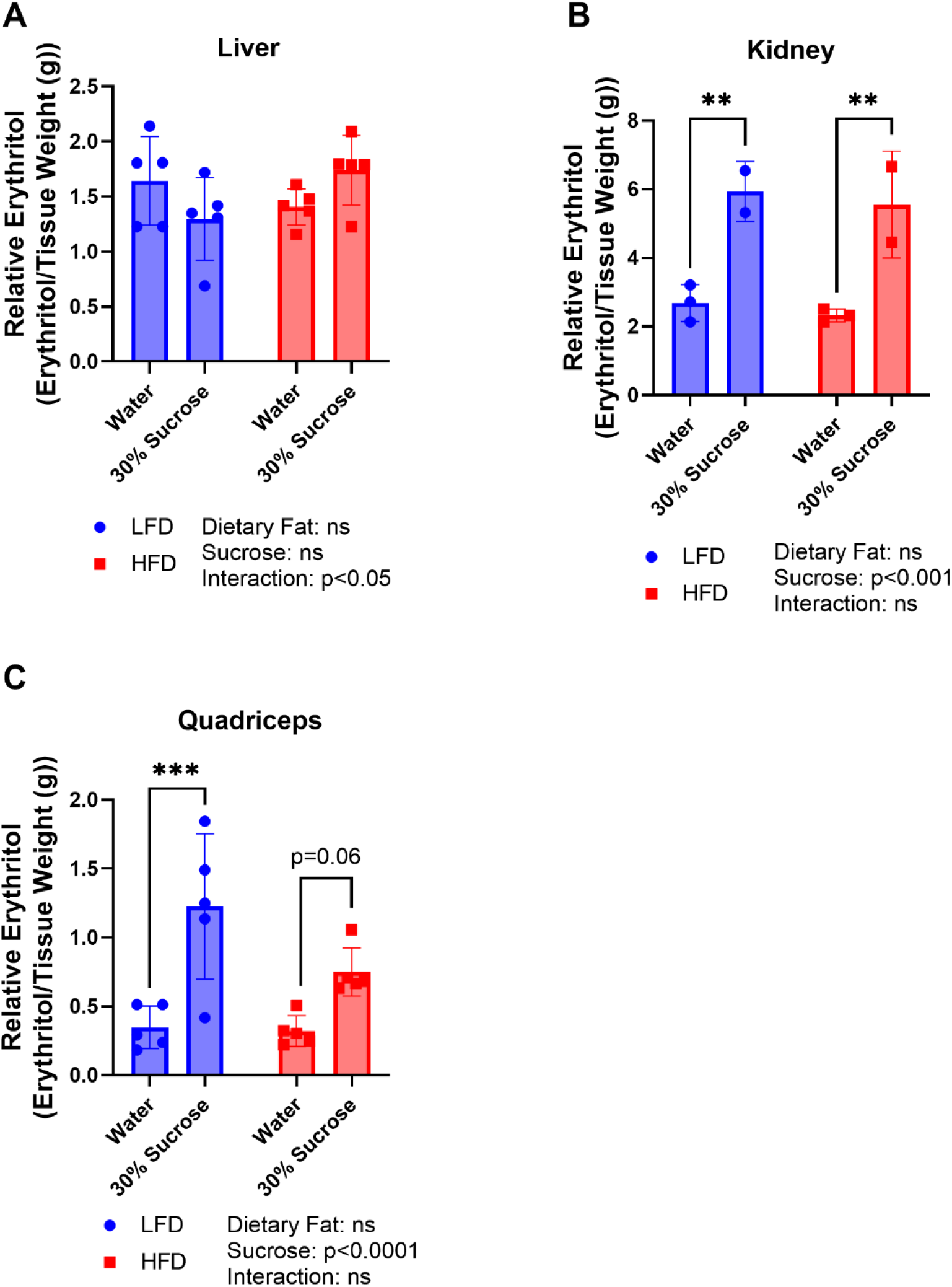
Sucrose significantly elevates erythritol in kidney and quadriceps. Relative erythritol content of A) liver, B) kidney, and C) quadriceps of mice following 8 weeks of dietary treatment. Data are presented as mean ± SD. **p<0.01, ***p<0.001. HFD: high-fat diet; LFD: low-fat diet.

### SORD deletion reduces tissue erythritol following exposure to sucrose water

Finally, we exposed SORD WT and KO animals to LFD with 30% sucrose for two weeks. We chose the SORD model based on the trend toward reduced fasted plasma erythritol on LFD and HFD (Fig. 2). We expected that increasing erythritol synthesis with dietary sucrose and measuring non-fasted plasma may amplify the effect of *Sord* loss. We found no difference in non-fasted plasma or urine erythritol between WT and KO mice **(Fig. 13A and 13B)**. In tissues, there was a significant effect of genotype and tissue-type on erythritol content (Fig. 13C, ANOVA main effect of genotype p<0.01, main effect of tissue p<0.0001). Notably, the kidneys of KO animals contained 30% less erythritol than WT controls (Fig. 13C p<0.05). There was no difference in quadriceps erythritol between genotypes. There was also no difference in body weight, food intake, or non-fasted blood glucose between genotypes **(Fig. S12A-S12C)**.

**Figure 13.**
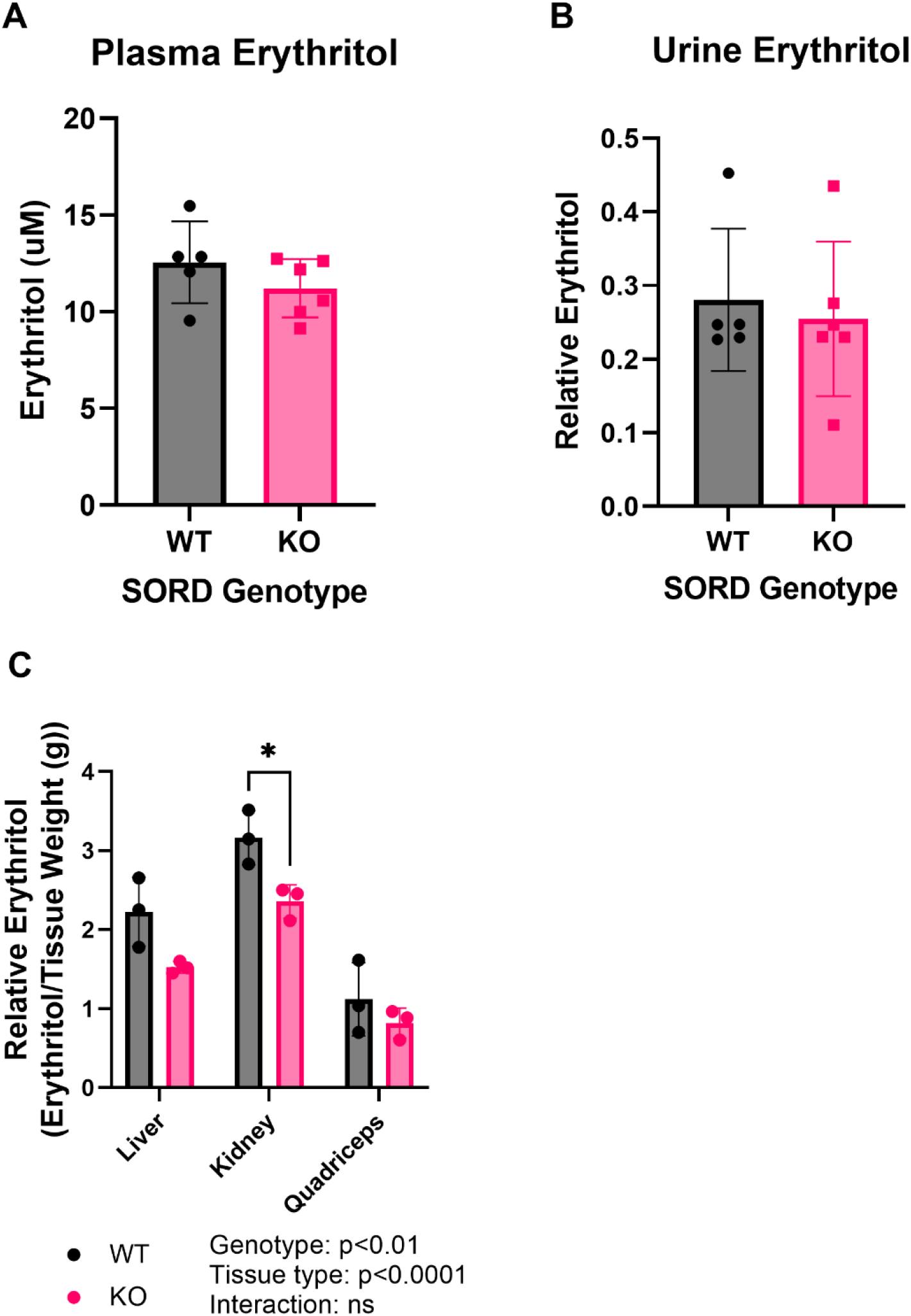
SORD null animals have reduced tissue erythritol in response to sucrose overfeeding. A) Non-fasted plasma erythritol, B) relative urine erythritol, and C) relative tissue erythritol in SORD WT and KO mice exposed to LFD with 30% sucrose. Data are presented as mean ± SD. *p<0.05. KO: knockout; WT: wildtype.

## Discussion

Surprisingly, we found no differences in plasma or tissue erythritol in mice lacking either *Sord* or *Adh1* expression. This was true in both diet-induced obese (HFD-fed mice) and LFD-fed mice (Figures 2-6), though there was a trend toward a reduction in plasma erythritol in *Sord*^*-/-*^ mice after 5 weeks (Figure 2B). There was also no evidence of compensation for SORD or ADH1 loss with an increase in protein levels of the alternative enzyme (Figures 4 and 7). This suggests that basal levels of ADH1 or SORD are sufficient to maintain erythritol synthesis when one is lost.

In cell culture models, SORD knockdown reduces erythritol synthesis only under high-glucose conditions (14). We hypothesized that SORD may also only be essential for erythritol synthesis when dietary sugar is in excess. To test this, we exposed SORD WT and KO mice to 30% sucrose in drinking water for two weeks, a relatively short exposure that did not result in effects on body weight (Fig S12A). There was no effect of *Sord* genotype on plasma or urine erythritol in response to sugar overfeeding (Figure 12). There was, however, a 30% reduction in kidney erythritol in SORD KO mice after sugar overfeeding.

Our findings also indicate that neither excess caloric intake from fat nor hyperglycemia are the driving factor in erythritol synthesis in young mice. Mice fed HFD exhibited elevated caloric intake, body weight gain, and fasting glucose compared to LFD controls with no significant difference in erythritol synthesis (Figures 8 and 10). In contrast, exposure to 30% sucrose in drinking water elevated non-fasted plasma and urine erythritol over the course of 8 weeks (Figures 10 and 11). Mice fed LFD with 30% sucrose water consistently exhibited the highest plasma and urine erythritol content (Figures 10 and 11). These mice consumed more calories from sugar than any other dietary treatment (Figure 8D). Mice fed HFD with 30% sugar water also exhibited elevated non-fasting plasma and urine erythritol compared to water controls, but lower plasma and urine erythritol levels than mice consuming LFD with 30% sucrose (Figures 10 and 11). The difference between these sucrose-exposure groups is likely due to the amount of sugar water consumed in the LFD group, rather than an interaction between dietary fat and dietary sugar. On average, mice fed LFD with 30% sucrose consumed 55% more calories from sugar water than mice fed HFD with 30% sucrose (Fig 8D).

Erythritol synthesis *in vivo* appears to be controlled by simple sugar consumption rather than total carbohydrate intake. This is supported by the comparison of plasma and urinary erythritol in LFD and HFD-fed mice. The LFD (10% FDC) contains more carbohydrates than the HFD (60% FDC). However, there is no difference in circulating erythritol levels between mice exposed to these diets (Figure 10). The primary carbohydrate source in the 10% FDC diet is cornstarch, which makes up 40% of the calories. Cornstarch is primarily composed of amylopectin, a branching, slowly digested chain of glucose that has a low glycemic index (18,19). Erythritol synthesis appears to respond to rapid (i.e. sucrose in drinking water) rather than slowly digestible carbohydrates. Other studies have shown differences between liquid and solid sucrose administration on the metabolic response to sugar in mice (20,21). Further studies are required to determine if erythritol synthesis is consistently elevated by simple, not complex carbohydrate intake when controlling for mode of administration (liquid/solid).

In humans, elevated erythritol in fasting plasma is a biomarker for cardiometabolic disease risk (1–3). Despite significant elevation in non-fasted circulating erythritol, we found no impact of diet on fasted plasma or urine erythritol levels (Figures 10C and 11C). There are several factors that may contribute to the lack of changes in fasting plasma erythritol in mouse models. This work was performed in young, healthy mice, whereas the human observational studies have largely been performed on middle-aged participants (2,3,5–8,10). These findings may also reflect species-specific differences in metabolism. Mice have an overall more rapid metabolism and higher glucose turnover compared to humans (22,23). In addition, fasting mice deplete available glucose more rapidly than humans and are more reliant on gluconeogenesis to maintain glucose homeostasis (22,23). The rapid depletion of glucose stores in fasting mice may limit the production of erythritol from glucose catabolism. To overcome the limitation of mouse models, future work in humans is required to determine if sugar consumption contributes to elevated fasting erythritol.

Finally, we found that sucrose water elevated kidney and quadriceps erythritol content, while liver erythritol content was stable across all diets (Figure 12). Differences in liver erythritol may have been dampened by lipid accumulation on sucrose diets. During tissue erythritol quantification, polar metabolites are normalized to tissue wet weight, which is elevated by lipid deposition and may reduce the relative erythritol content per gram of tissue. Overall, however, it appears that the liver is not driving sucrose-induced erythritol synthesis. Elevated kidney erythritol was an expected response based on previous work in human proximal tubule cells in which high glucose media increased intracellular erythritol (14,24). Elevated circulating erythritol is also associated with markers of impaired kidney function (11,25).

This is the first report of elevated erythritol in skeletal muscle in response to glucose availability (Figure 12C). This was unexpected based on the low expression and activity of pentose phosphate pathway enzymes in skeletal muscle and the relatively low quadriceps erythritol content compared to liver and kidneys (13,26,27). In humans, however, Lustgarten and Fielding recently observed a negative association between skeletal muscle density and serum erythritol levels (25). Pentose phosphate pathway enzymes have also been shown to increase in mouse skeletal muscle in response to high-fat diet exposure, and muscle glucose-6-phosphate dehydrogenase (G6PD) expression is associated with impaired glucose metabolism (26,28,29). Historically, increased PPP activity has been reported following skeletal muscle injury and in disordered skeletal muscle (30,31). Taken together, these findings suggestion that circulating erythritol may be indicative of impaired skeletal muscle metabolism.

The skeletal muscle may also be a useful model for further understanding the regulation of erythritol synthesis. Although quadriceps appear to have lower erythritol content compared to the liver and kidneys, this represents a small percentage of total body skeletal muscle (13). Skeletal muscle is responsible for around 30% of postprandial glucose disposal (whereas the kidney disposes of only 7%) and may therefore contribute more to circulating erythritol levels than is captured by a single tissue sample (22). It is notable that the erythritol-synthesizing enzymes ADH1 and SORD were originally identified and purified from rabbit liver (12). Muscle ADH1 and SORD protein levels are relatively low, which makes it an ideal tissue to identify alternative enzymes that catalyze the conversion of erythrose to erythritol (32). The modest expression of ADH1 and SORD in skeletal muscle may also have blunted the effect of *Adh1*/*Sord* knockout on plasma erythritol levels.

A limitation of this study is that protein intake was not constant between the 30% sucrose and the water control groups. *Ad libidum* access to sucrose solution reduced solid food intake in both LFD with 30% sucrose and HFD with 30% sucrose mice, resulting in a decrease in protein intake. Future studies may be able to account for protein intake with pair feeding, which is beyond the scope of this work. Circulating erythritol was proportional to the amount of sucrose water consumed, suggesting that sucrose was the primary determinant of elevated erythritol levels in the present study.

In conclusion, we found that sucrose intake significantly elevated erythritol synthesis and excretion in mice. Erythritol synthesis and excretion is a novel pathway for the disposal of glucose carbons when dietary sugar is in excess. Future studies in humans should assess if there is an association between simple sugar intake and erythritol in plasma and/or skeletal muscle.

## Abbreviations

ADH1: alcohol dehydrogenase 1
DIO: diet-induced obesity
G6PD: glucose-6-phosphate dehydrogenase
HFD: high-fat diet
IPGTT: intraperitoneal glucose tolerance test
KO: knockout
LFD: low-fat diet
PPP: pentose phosphate pathway
SORD: sorbitol dehydrogenase
TKT: transketolase
WT: wildtype.

## Acknowledgements

The authors thank Peyton Carpen and Brian Walker for assistance with data collection and the Cornell Statistical Consulting Unit for assistance with data analysis.

## Author Contributions

SRO and MSF designed research; SRO conducted research and analyzed data; SRO and MSF wrote the paper. MSF had primary responsibility for final content. All authors have read and approved the final manuscript.

