## supplemental figures for "Elevated plasma and urinary erythritol is a biomarker of excess simple carbohydrate intake in mice"

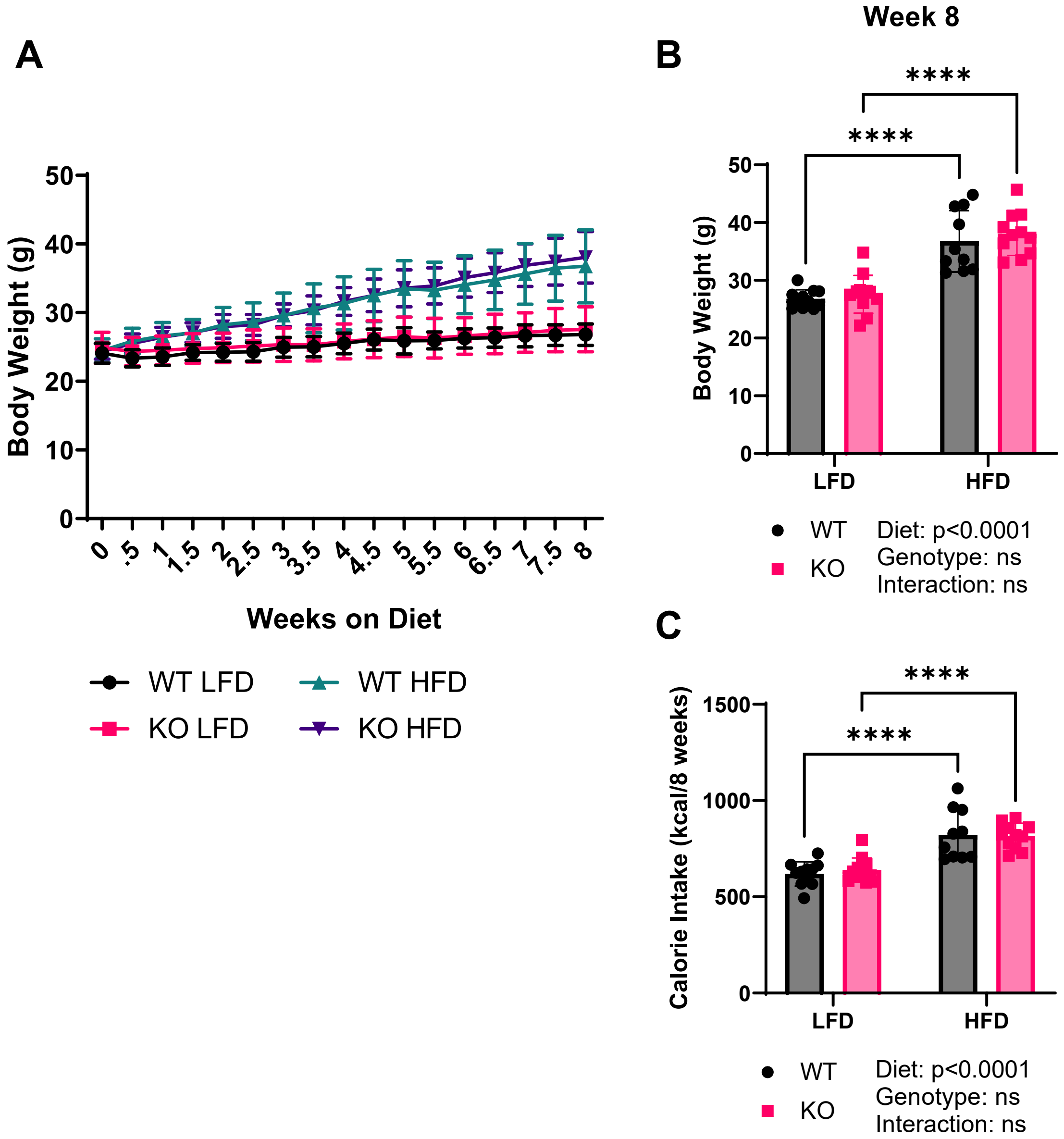


**Figure S1. Loss of SORD does not impact body weight or caloric intake.** A) Body weight in grams over time. B) Body weight in grams following 8 weeks of dietary treatment. C) Total cumulative caloric intake in kilocalories (kcal). Data are expressed as mean ± SD. ****p<0.0001. HFD: high-fat diet; KO: knockout; LFD: low-fat diet; WT: wildtype.

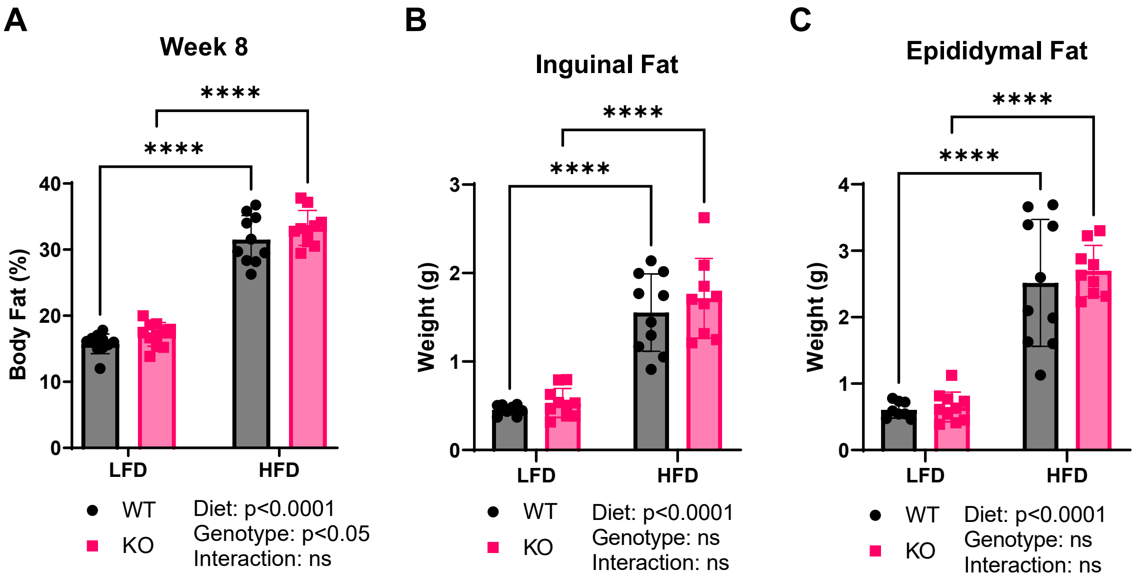


**Figure S2. SORD knockout does not impact total body fat percentage or adipose depot weight compared to WT littermates.** A) Body fat percentage, measured by NMR, B) inguinal fat weight in grams, and C) epididymal fat weight in grams after 8 weeks on experimental diets. Data are expressed as mean ± SD. ****p<0.0001. HFD: high-fat diet; KO: knockout; LFD: low-fat diet; WT: wildtype.

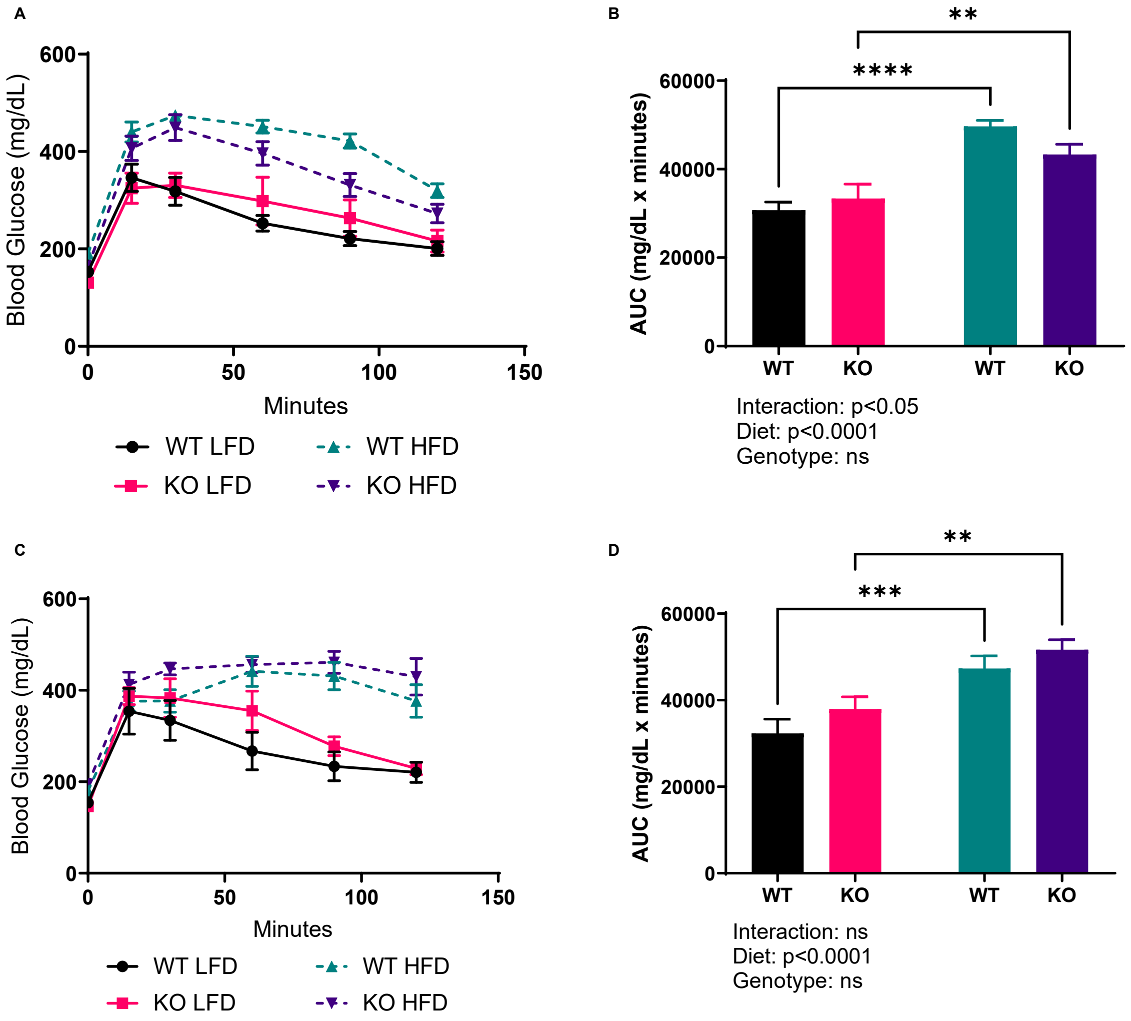


**Figure S3. Loss of SORD does not impact glucose tolerance.** A) Results of intraperitoneal glucose tolerance testing (IPGTT) and B) area under the curve (AUC) of IPGTT after 2 weeks on experimental diets. C) IPGTT results after 8 weeks on experimental diets and D) the AUC of panel C. Panels A and C are presented as mean ± SEM, n=5, and panels B and D are presented as mean ± SD. **p<0.01, ***p<0.001. AUC: area under the curve; HFD: high-fat diet; KO: knockout; LFD: low-fat diet; WT: wildtype.

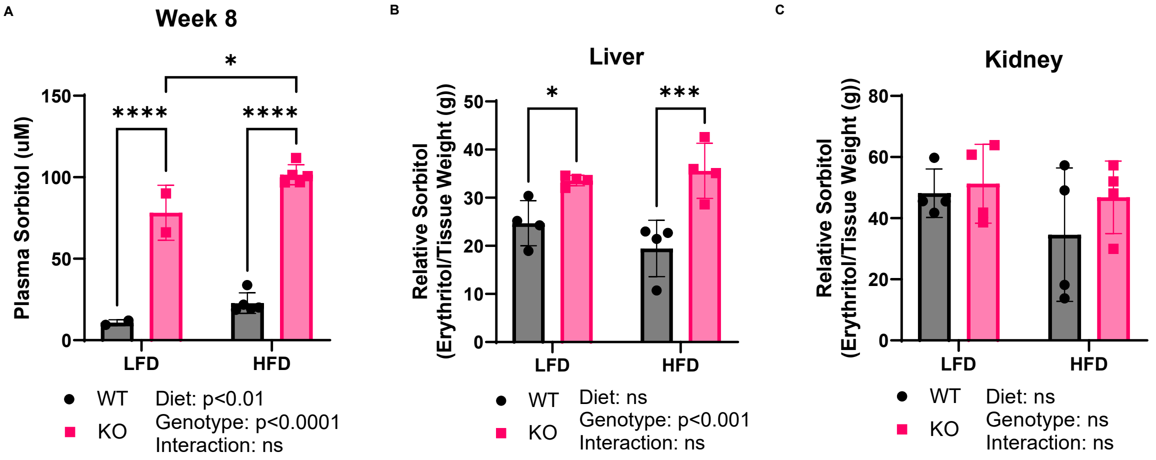


**Figure S4. Loss of SORD significantly elevates plasma and liver sorbitol.** A) Plasma sorbitol in SORD WT and KO mice following 8 weeks of exposure to LFD or HFD. B) Relative tissue sorbitol in liver and C) kidney in WT and KO mice after 8 weeks of dietary treatment. Data are shown as mean ± SD. $^*p<0.05$, $^{***}p<0.001$, $^{****}p<0.0001$. HFD: high-fat diet; KO: knockout; LFD: low-fat diet; WT: wildtype.

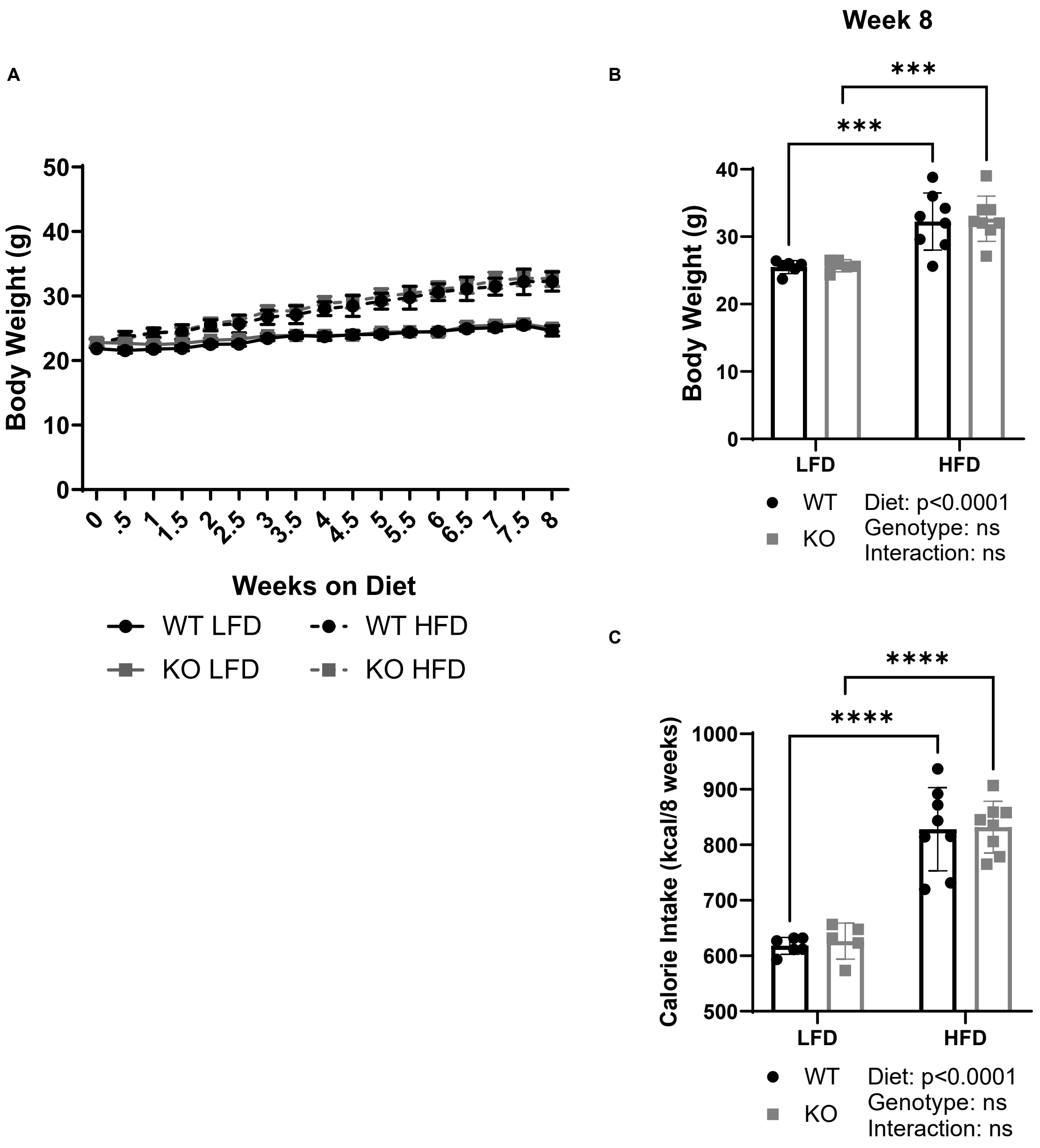


**Figure S5. ADH1 deletion does not impact body weight or caloric intake.** A) Body weight in grams over time. B) Body weight in grams following 8 weeks of dietary treatment. C) Total cumulative caloric intake in kilocalories (kcal). Data are expressed as mean ± SD. ***$p<0.001$, ****$p<0.0001$. HFD: high-fat diet; KO: knockout; LFD: low-fat diet; WT: wildtype.

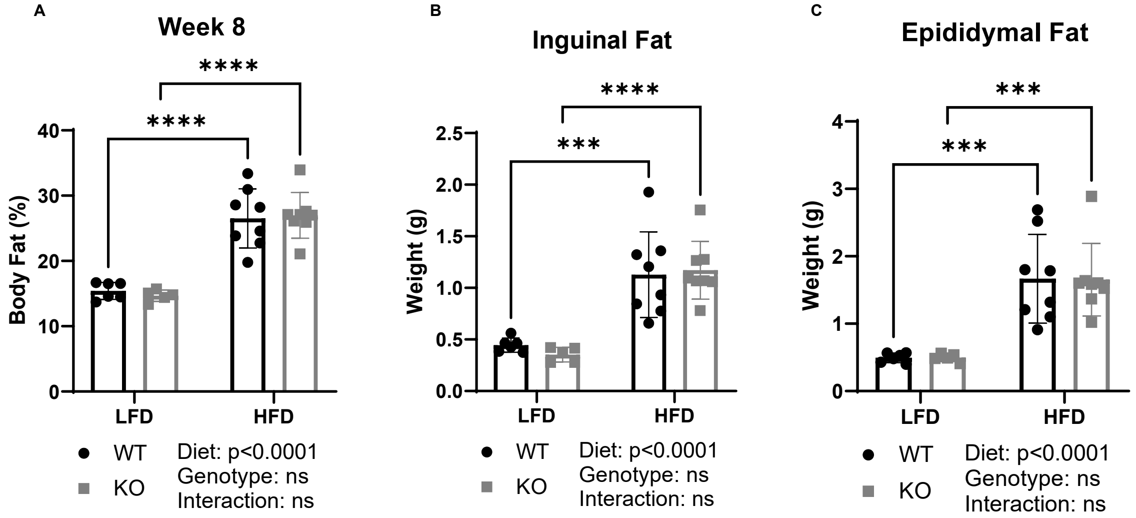


**Figure S6. Loss of ADH1 does not significantly impact total body fat percentage.** A) Body fat percentage, measured by NMR, B) inguinal fat weight in grams, and C) epididymal fat weight in grams after 8 weeks on experimental diets. Data are expressed as mean ± SD. ***p<0.001 ****p<0.0001. HFD: high-fat diet; KO: knockout; LFD: low-fat diet; WT: wildtype.

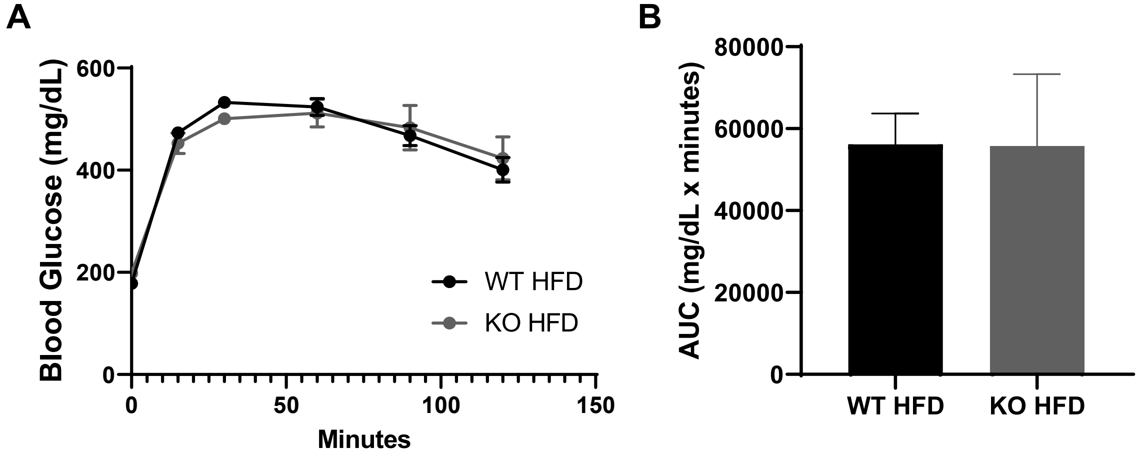


**Figure S7. ADH1 knockout mice fed HFD do not have altered glucose tolerance compared to WT littermates.** A) Results of intraperitoneal glucose tolerance testing (IPGTT) and B) area under the curve (AUC) of IPGTT after 8 weeks on experimental diets. Panels A is presented as mean ± SEM, n=5, and panels B is mean ± SD. AUC: area under the curve; HFD: high-fat diet; KO: knockout; WT: wildtype.

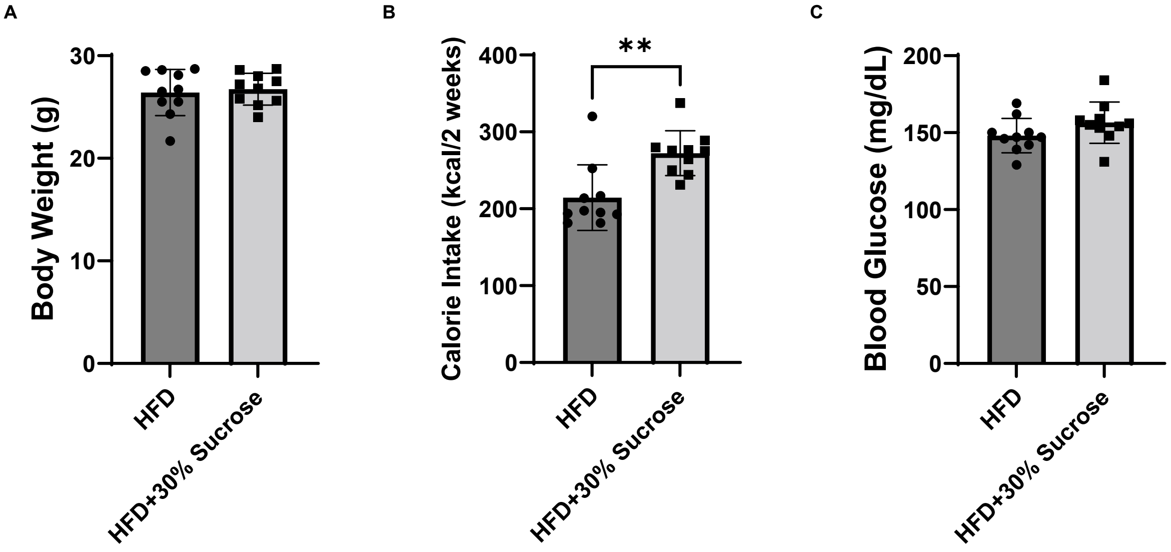


**Figure S8. Sucrose water increases caloric intake but does not impact body weight or blood glucose levels.** A) Body weight in grams following 2 weeks of treatment with water or 30% sucrose on HFD. B) Total caloric intake over 2 weeks of water treatment. C) Non-fasted blood glucose following 2 weeks of water or sucrose treatment. Data are shown as mean ± SD. **p<0.01. HFD: high-fat diet.

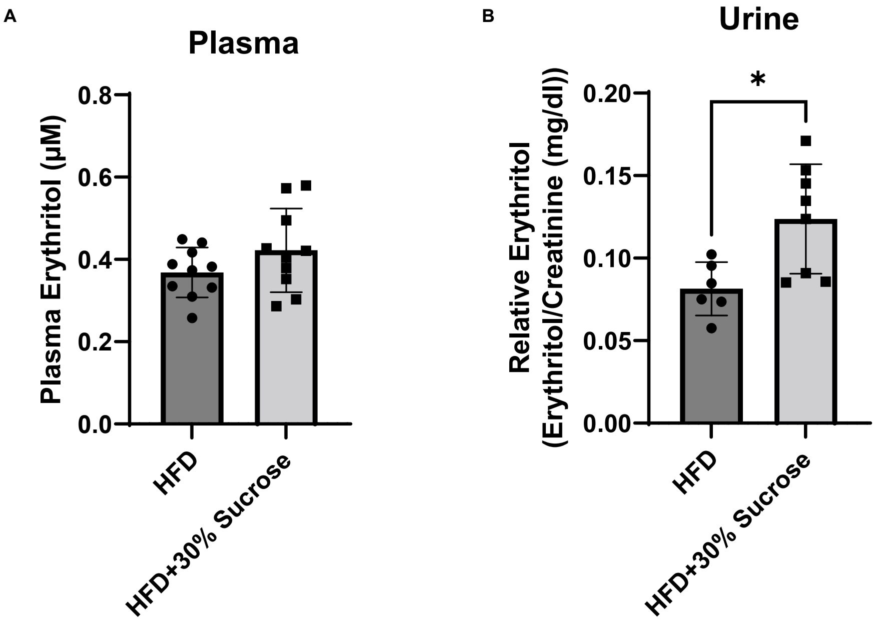


**Figure S9. Two-week exposure to sucrose water elevates urine erythritol.** A) Fasted plasma erythritol and B) non-fasted relative erythritol in urine after 2 weeks of water or 30% sucrose exposure. Data presented as mean ± SD. *p<0.05. HFD: high-fat diet.

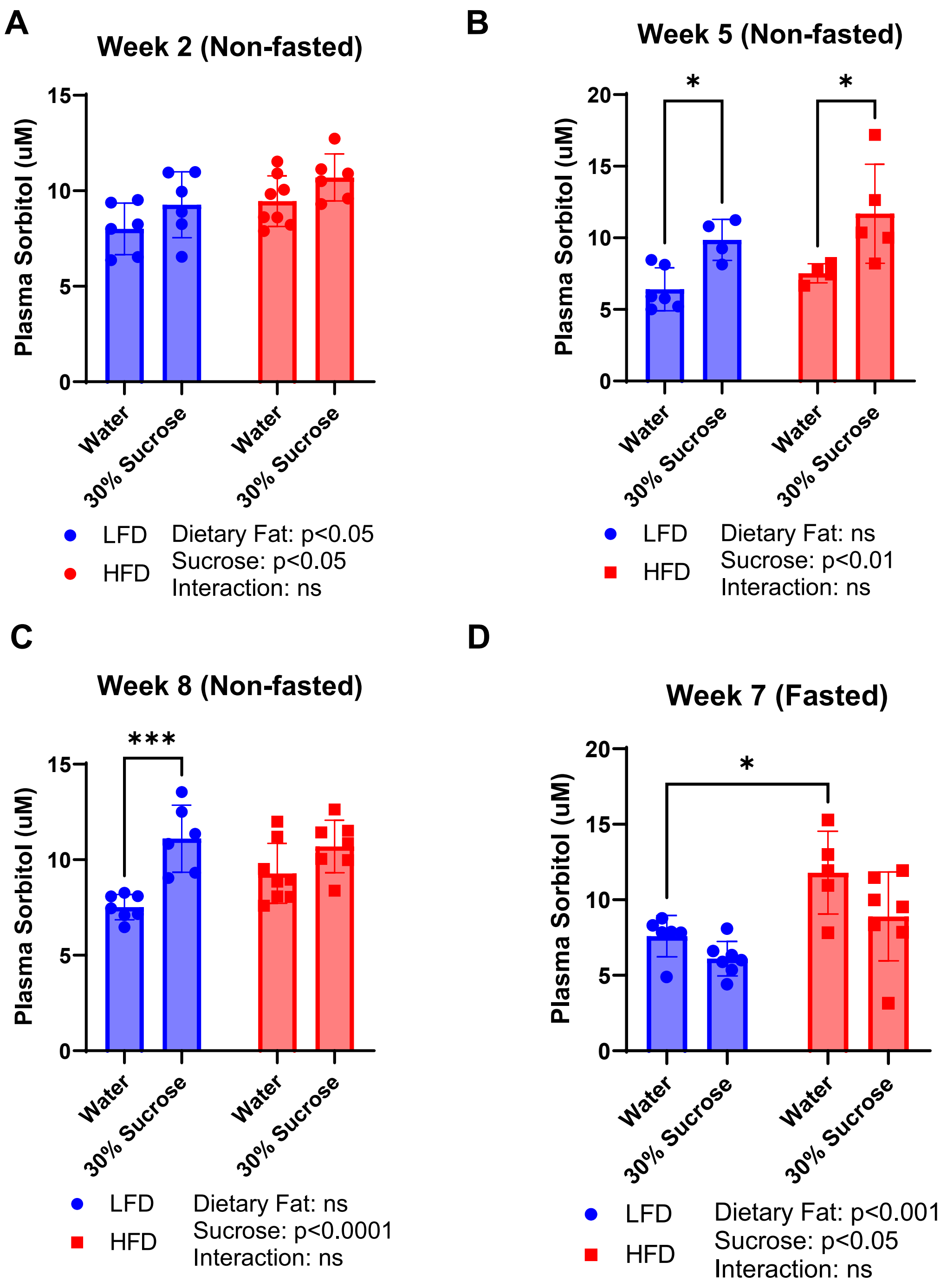


**Figure S10. Response of plasma sorbitol to diet composition differs between fed and fasted mice.** Plasma sorbitol in non-fasted mice at A) 2 weeks, B) 5 weeks, and C) 8 weeks of diet treatment. D) Plasma sorbitol in fasted mice after 7 weeks of dietary treatment. Data are presented as mean ± SD. *p<0.05, ***p<0.001. HFD: high-fat diet; LFD: low-fat diet.

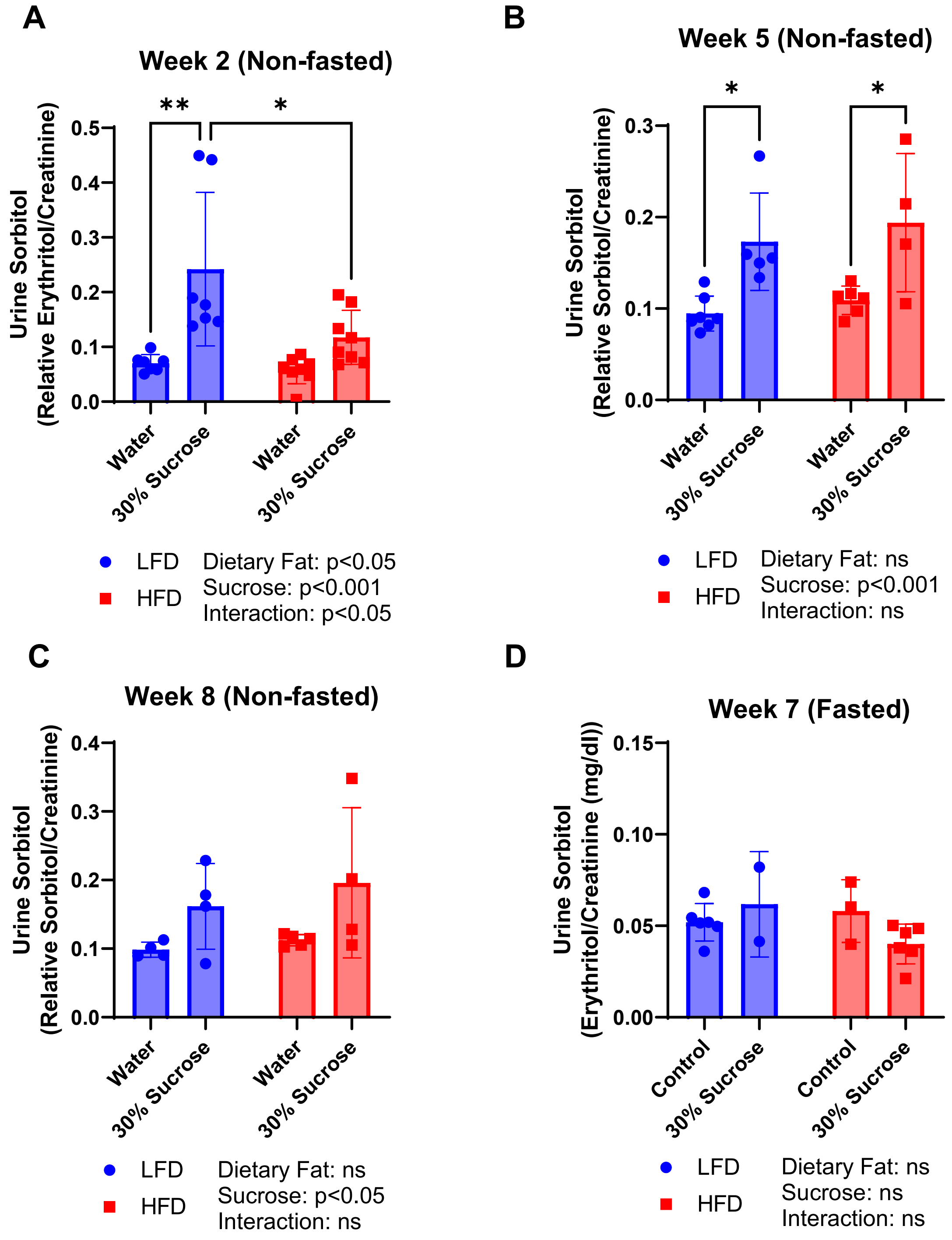


**Figure S11. Urinary sorbitol is elevated by sucrose in drinking water in fed mice.** Relative urinary sorbitol in non-fasted mice at A) 2 weeks, B) 5 weeks, and C) 8 weeks of diet treatment. D) Relative urinary sorbitol in fasted mice after 7 weeks of dietary treatment. Sorbitol was normalized to urinary creatinine content. Data are presented as mean ± SD. *p<0.05, **p<0.01. HFD: high-fat diet; LFD: low-fat diet.

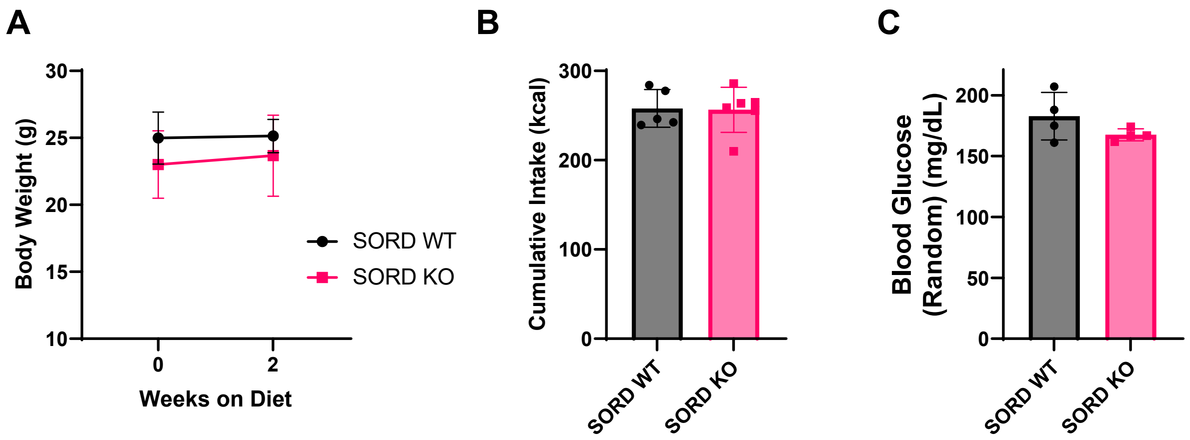


**Figure S12. SORD knockout does not impact body weight, food intake, or random blood glucose in response to sugar water.** A) Body weight in grams over time in SORD WT and KO mice fed LFD with 30% sucrose. B) Total caloric intake in kilocalories (kcal) over 2 weeks. C) Non-fasted blood glucose after 2 weeks exposure to LFD with 30% sucrose. Data expressed as mean ± SD. KO: knockout; LFD: low-fat diet; SORD: sorbitol dehydrogenase; WT: wildtype.
