## supplemental tables for "Elevated plasma and urinary erythritol is a biomarker of excess simple carbohydrate intake in mice"

**Online supporting material** for Ortiz, S.R, “Elevated plasma and urinary erythritol is a biomarker of excess simple carbohydrate in mice”

| DYET# 104783 Custom Low Fat (10% FDC) AIN-93G Based Diet | | | | |
|---|---|---|---|---|
| **Ingredient** | **kcal/gm** | **grams/kg** | **kcal/kg** | **%kcal** |
| Sterile Casein | 3.72 | 200 | 744 | 20.47274 |
| L-Cystine | 4 | 3 | 12 | 0.330206 |
| Sucrose | 4 | 100 | 400 | 11.00685 |
| Cornstarch | 3.6 | 413.496 | 1488.6 | 40.962 |
| Dyetrose | 3.8 | 145 | 551 | 15.16194 |
| Soybean Oil | 9 | 20 | 180 | 4.953083 |
| t-Butylhydroquinone | 0 | 0.004 | 0 | 0 |
| Lard | 9 | 21 | 189 | 5.200737 |
| Cellulose | 0 | 50 | 0 | 0 |
| Mineral Mix #210025 | 0.88 | 35 | 30.8 | 0.847528 |
| Vitamin Mix #310025 | 3.87 | 10 | 38.7 | 1.064913 |
| Choline Bitartrate | 0 | 2.5 | 0 | 0 |

**Supplementary Table 1. Composition of low-fat diet (LFD).** Diets were obtained from Dyets Inc., Bethlehem PA. FDC: fat-derived calories; LFD: low-fat diet.

| DYET# 103651 Dyets Version of AIN-93G Based Diet Induced Obesity (DIO) Diet with 60% Fat Derived Calories | | | | |
|---|---|---|---|---|
| **Ingredient** | **kcal/gm** | **grams/kg** | **kcal/kg** | **%kcal** |
| Sterile Casein | 3.72 | 265 | 985.8 | 19.12954 |
| L-Cystine | 4 | 4 | 16 | 0.310481 |
| Sucrose | 4 | 90 | 360 | 6.985832 |
| Maltose dextrin | 3.8 | 159.994 | 607.98 | 11.79791 |
| Soybean Oil | 9 | 30 | 270 | 5.239374 |
| t-Butylhydroquinone | 0 | 0.006 | 0 | 0 |
| Lard | 9 | 310 | 2790 | 54.1402 |
| Cellulose | 0 | 65.5 | 0 | 0 |
| Mineral Mix #210025 | 0.88 | 48 | 42.24 | 0.819671 |
| Vitamin Mix #310025 | 3.87 | 21 | 81.27 | 1.577052 |
| Choline Bitartrate | 0 | 2.5 | 0 | 0 |

**Supplementary Table 2. Composition of high-fat diet (HFD).** Diets were obtained from Dyets Inc., Bethlehem PA. FDC: fat-derived calories; HFD: high-fat diet.
